# Molecular Insights into the Production of Extracellular Vesicles by Plants

**DOI:** 10.1101/2025.06.16.659989

**Authors:** Benjamin L. Koch, Dillon Gardner, Hannah Smith, Rachel Bracewell, Linnaea Awdey, Jessica Foster, M. Lucía Borniego, David H. Munch, Mads E. Nielsen, Raghavendra Pasupuleti, Jonathan Trinidad, Brian Rutter, Hans Thordal-Christensen, Roger W. Innes

## Abstract

Extracellular vesicles (EVs) produced by Arabidopsis are highly heterogeneous in protein content. Specific EV subpopulations have been proposed to participate in plant immunity, particularly during plant-fungal interaction. To understand the origins of plant EV heterogeneity, we used a proximity labeling approach to identify proteins and pathways involved in the secretion of distinct EV subpopulations. Proximity labeling, co-immunoprecipitation, and fluorescence microscopy in *Nicotiana benthamiana* all indicated a general role in EV secretion for EXO70 proteins (a subunit of the exocyst complex) and the immune-related protein RPM1-INTERACTING PROTEIN 4 (RIN4), while the ER-localized VAMP-ASSOCIATED PROTEIN 27(VAP27) was specifically associated with the EV marker protein TETRAPSANIN8 (TET8). Despite being secreted in separate EV populations, we found that TET8 and PENETRATION 1 (PEN1) co-localized in multivesicular body-like subcellular structures. Based on these results, we tested Arabidopsis mutants and identified several proteins involved in EV secretion, including members of the *exo70* family, *rin4*, *rabA2a*, *scd1* (a GTP-exchange factor for RabE GTPases), and *vap27*. We also uncovered a possible connection between trichomes and EVs, as the trichomeless mutant *glabrous1* secreted approximately half the number of EVs as wild type. Furthermore, PEN1 MVB-like structures were prevalent in guard cells, suggesting that guard cells may contribute to secretion of PEN1+ EVs on the leaf surface. Lastly, we found that *exo70* family mutants are more susceptible to infection with the fungal pathogen *Colletotrichum higginsianum*, underlining the importance of secretion for plant immunity. Together, our results unravel some of the complex mechanisms that give rise to EV subpopulations in plants.

## Introduction

Extracellular vesicles (EVs) are secreted by organisms from all three domains of life (Gill et al., 2019). EVs range in size from 50 nm to 5 microns in diameter and can encapsulate or associate with proteins, nucleic acids, metabolites, and other molecules. In mammals, EVs have been shown to contribute to antigen presentation as well as delivery of cargo to recipient cells (Raposo et al., 1996; Verweij et al., 2019), but EVs can play a myriad of roles depending on the needs of their organism, such as in plasma membrane repair (Williams et al., 2023), nutrient homeostasis (Biller et al., 2014; Gorlas et al., 2015), as viral decoys (Biller et al., 2014), in biofilm formation for drug resistance (Zarnowski et al., 2018; Johnston et al., 2023), as vectors for plasmid delivery (Erdmann et al., 2017), and for cross-kingdom communication (Micali et al., 2011; Cai et al., 2018; He et al., 2021). The pool of EVs secreted by a given organism is comprised of multiple subpopulations of EVs distinguishable by their fundamental characteristics such as size, density, and cargo, which may enable them to perform distinct functions (Willms et al., 2016; Tkach et al., 2017; Jeppesen et al., 2019).

In plants, EVs of the model organism Arabidopsis contain stress-response proteins and EV secretion is induced during biotic and abiotic stress (Regente et al., 2017; Rutter and Innes, 2017; Chaya et al., 2024; Koch et al., 2025). Plant EVs have long been linked with pathogen infection by ultrastructural analysis, and plant EVs and EV marker proteins focally accumulate at the site of infection (Ehrlich et al., 1968; Collins et al., 2003; Assaad et al., 2004; An et al., 2006; Meyer et al., 2009; Micali et al., 2011; Rubiato et al., 2022). EVs have been proposed to contribute to pre-penetration resistance, papillae formation, transfer of nucleic acids for cross-kingdom communication, and other antimicrobial functions (Regente et al., 2017; Cai et al., 2018; He et al., 2021; Wang et al., 2024). EV-associated proteins may contribute to these functions, or to earlier steps in EV formation. For instance, the syntaxin protein PEN1 localizes to both the plasma membrane and to EVs and contributes to papillae formation, likely by promoting membrane fusion events associated with formation of EVs (Nielsen et al., 2012; Rubiato et al., 2022). Consistent with EVs contributing to multiple functions, Arabidopsis secretes diverse subpopulations of EVs (Koch et al., 2025). Different isolation and purification techniques can be used to partially separate Arabidopsis EV subpopulations, and total internal fluorescence (TIRF) microscopy has been used to directly observe distinct subpopulations, particularly subpopulations marked by the syntaxin protein PENETRATION1 (PEN1; also known as SYP121) and TETRASPANIN8 (TET8) (He et al., 2021; Ghosh and Innes, 2025; Koch et al., 2025). Plant EV subpopulations respond differentially to biotic and abiotic stress, with TET8, PEN1, and RPM1-interactin protein 4 (RIN4) marking particularly responsive EV subpopulations (Rutter and Innes, 2017; Koch et al., 2025).

In other organisms, the diversity of EVs is likely the result of complex mechanisms that control EV biogenesis. Mammalian cells secrete three basic classes of EVs: microvesicles, which bud directly from the plasma membrane (50-1000nm), apoptotic bodies, which are released upon cell death (100-5000nm), and exosomes, which are secreted through the fusion of a multivesicular body (MVB) with the plasma membrane (50-200nm) (Jeppesen et al., 2019; Jeppesen et al., 2023). Exosome biogenesis involves the assembly of endosomal sorting complexes required for transport (ESCRT) proteins at the surface of MVBs, which assist in creating intraluminal vesicles (ILVs) and loading of ILVs with cargo proteins (Babst, 1998; Katzmann et al., 2001; Babst et al., 2002; Raiborg et al., 2002; Bache et al., 2003; Lu et al., 2003; Muzioł et al., 2006; Raiborg and Stenmark, 2009). In human cells, knockdown of ESCRTs or ESCRT-associated proteins reduces the secretion of multiple EV subpopulations (Baietti et al., 2012; Colombo et al., 2013). ESCRT components are largely conserved among eukaryotes and archaea, and knockdown of ESCRTs in the model yeast *Saccharomyces cerevisiae* (the organism in which ESCRTs were originally identified), the pathogenic fungus *Candida albicans*, and the hyperthermophilic and acidophilic archaeon *Sulfolobus islandicus* reduces EV secretion (Oliveira et al., 2010; Zarnowski et al., 2018; Zhao et al., 2019; Liu et al., 2021). However, ILVs can still form in the absence of ESCRTs (Stuffers et al., 2009). In animal cells, ESCRT-independent EV biogenesis can involve lipid modifying proteins, endoplasmic reticulum (ER) contact sites, liquid-liquid phase separation, and other mechanisms (Trajkovic et al., 2008; Larios et al., 2020; Liu et al., 2021; Barman et al., 2022). After ILV formation, the MVB can be directed with the help of Rab GTPases to the plasma membrane for secretion, sometimes in a subpopulation specific manner (Bobrie et al., 2012; Russell et al., 2012; Temoche-Diaz et al., 2019). On the way to the plasma membrane, MVBs can also interact with autophagosomes to form hybrid organelles, facilitating the secretion of autophagic substrates that would typically be bound for degradation (Guo et al., 2017; Jeppesen et al., 2019; Leidal et al., 2020; Peng et al., 2021). The biogenesis of microvesicles likely involves localization of proteins to microdomains in the plasma membrane, which are then released as EVs through a direct budding process (Nabhan et al., 2012; Nager et al., 2017; Beer et al., 2018). Annexins, calcium-regulated membrane-binding proteins, were thought to be markers of microvesicles in animal EVs, but some annexins can also mark MVB-derived EVs (Jeppesen et al., 2019; Williams et al., 2023).

Although progress is being made on the origins of animal EVs, very little is known about plant EV biogenesis. A recent study showed that TET8 may function as a carrier of <u>g</u>lycosylinositol<u>p</u>hospho-<u>c</u>eramides (GIPCs), a numerically dominant lipid of plant EVs, and that *tet8* mutant plants secrete fewer EVs, possibly due to altered Golgi morphology (the site of GIPC synthesis) and fewer ILVs per MVB (Liu et al., 2020; Liu et al., 2024).

To delve deeper into plant EV biogenesis, we employed proximity labeling to identify new proteins and pathways involved in the secretion of Arabidopsis EV subpopulations. Proximity labeling and follow-up experiments identified a general role in EV secretion for the EXO70 subunit of the exocyst complex, an octomeric tethering complex that is conserved among eukaryotes. Specific roles for VAMP-associated proteins (VAPs) and RabA2a were identified for TET8+ and PEN1+ EVs, respectively. Additionally, we identified a possible role for trichomes in EV secretion, though further investigation is needed to test whether trichomes are a significant source of EV secretion. Lastly, exocyst mutant plants were more susceptible to infection with the fungal pathogen *C. higginsianum*, highlighting the importance of secretion in pathogen resistance. This study lays the groundwork for future studies of plant EV biogenesis.

## Results

### Isolation of EVs from *Nicotiana benthamiana* leaves

We sought to determine the level of similarity between EV secretion systems of Arabidopsis and *N. benthamiana*. *N. benthamiana* EVs have been isolated and purified from apoplastic wash fluid (AWF) using protocols similar to Arabidopsis EV isolation (Zhang et al., 2020) (Ghosh and Innes, 2025). BLAST searches revealed that the *N. benthamiana* genome encodes orthologs of two well-characterized Arabidopsis EV marker proteins (Supplementary Fig. S1A). We thus tested antibodies raised to Arabidopsis EV marker proteins PEN1, RIN4, ANNEXIN2 (ANN2), and PATELLIN1 (PATL1). These four antibodies cross-reacted with proteins in *N. benthamiana* cell extracts (Supplementary Fig. S1B). We extracted apoplastic wash fluid from *N. benthamiana* leaves and pelleted at sequentially higher speeds (Supplementary Fig. S2B). Nanoparticle tracking analysis (NTA) showed that the majority of particles were present in the pellet at 40,000 × *g* (P40). These particles had typical EV morphology when imaged using negative stain TEM (Fig. 1A). Arabidopsis EV antibodies also recognized proteins in the *N. benthamiana* extracellular pellets, suggesting that the *N. benthamiana* orthologs of PEN1, RIN4, ANN2, and PATL1 could be potential markers of *N. benthamiana* EVs (Fig. 1A). ANN2, PATL1, and Rubisco Large Subunit (RBCL, the prominent band in Ponceau S) did not fully pellet in the P40 and remained in the supernatant (S100) even after the P100 spin, suggesting that these proteins could be at least partially secreted independent of EVs. Collectively, these observations confirmed that EVs can be isolated from *N. benthamiana* leaves using the protocol developed for Arabidopsis, and that EV cargo are conserved between Arabidopsis and *N. benthamiana*.

**Supplementary Figure S1.**
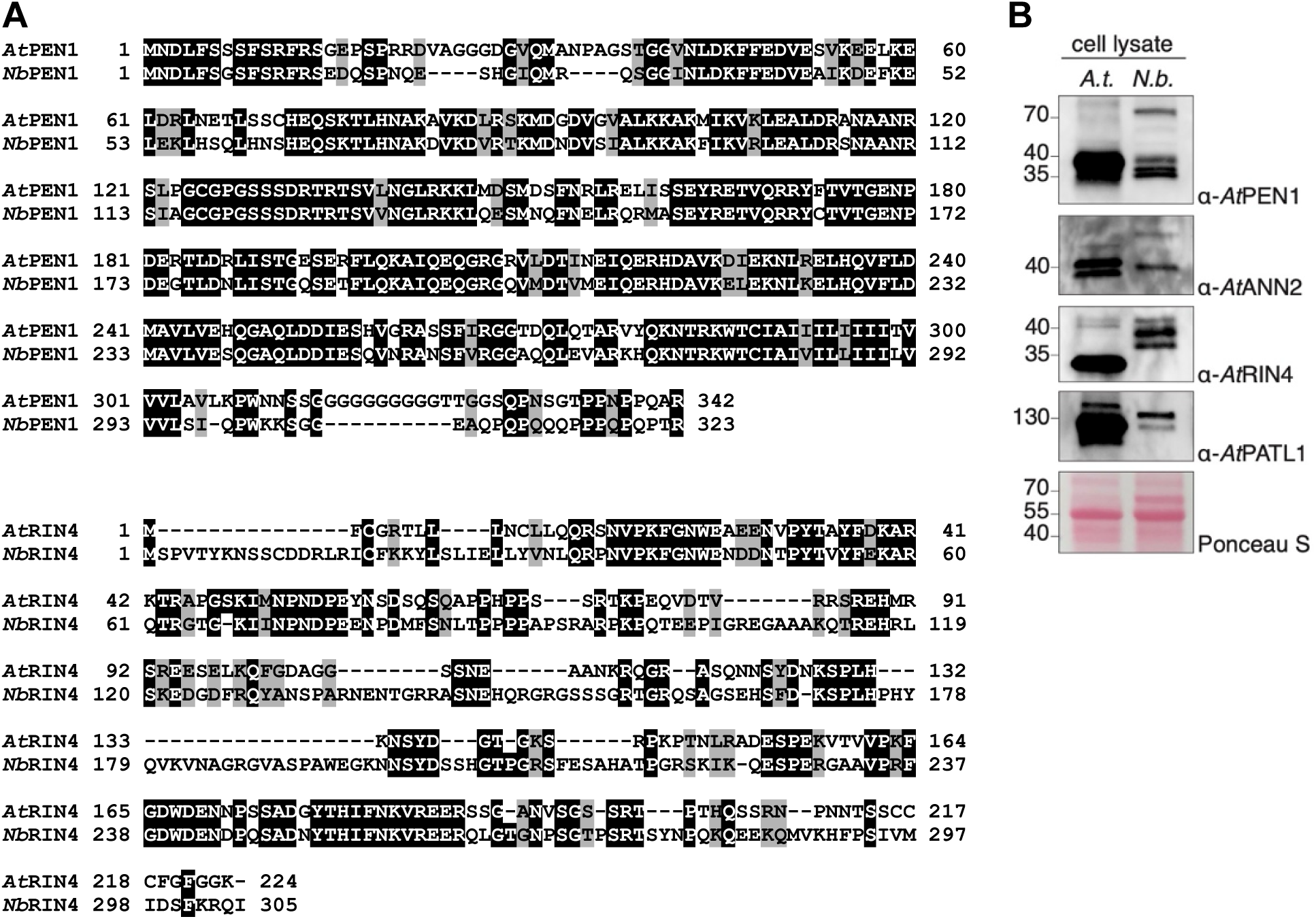
The Arabidopsis extracellular vesicle proteins PEN1 and RIN4 have orthologs in *N. benthamiana.* **A**) Sequence alignment of Arabidopsis and *N. benthamiana* PEN1 and RIN4 proteins. *N. benthamiana* orthologs were identified by performing a BLAST search of the *N. benthamiana* genome available at NbenBase (Kurotani et al., 2025). **B**) Equal amounts of leaf lysate from Arabidopsis (*A.t.*) and *N. benthamiana (N.b.)* were separated by SDS-PAGE and probed with antibodies raised to Arabidopsis EV marker proteins. Bands likely representing closely-related proteins were identified in *N. benthamiana* extracts.

**Figure 1.**
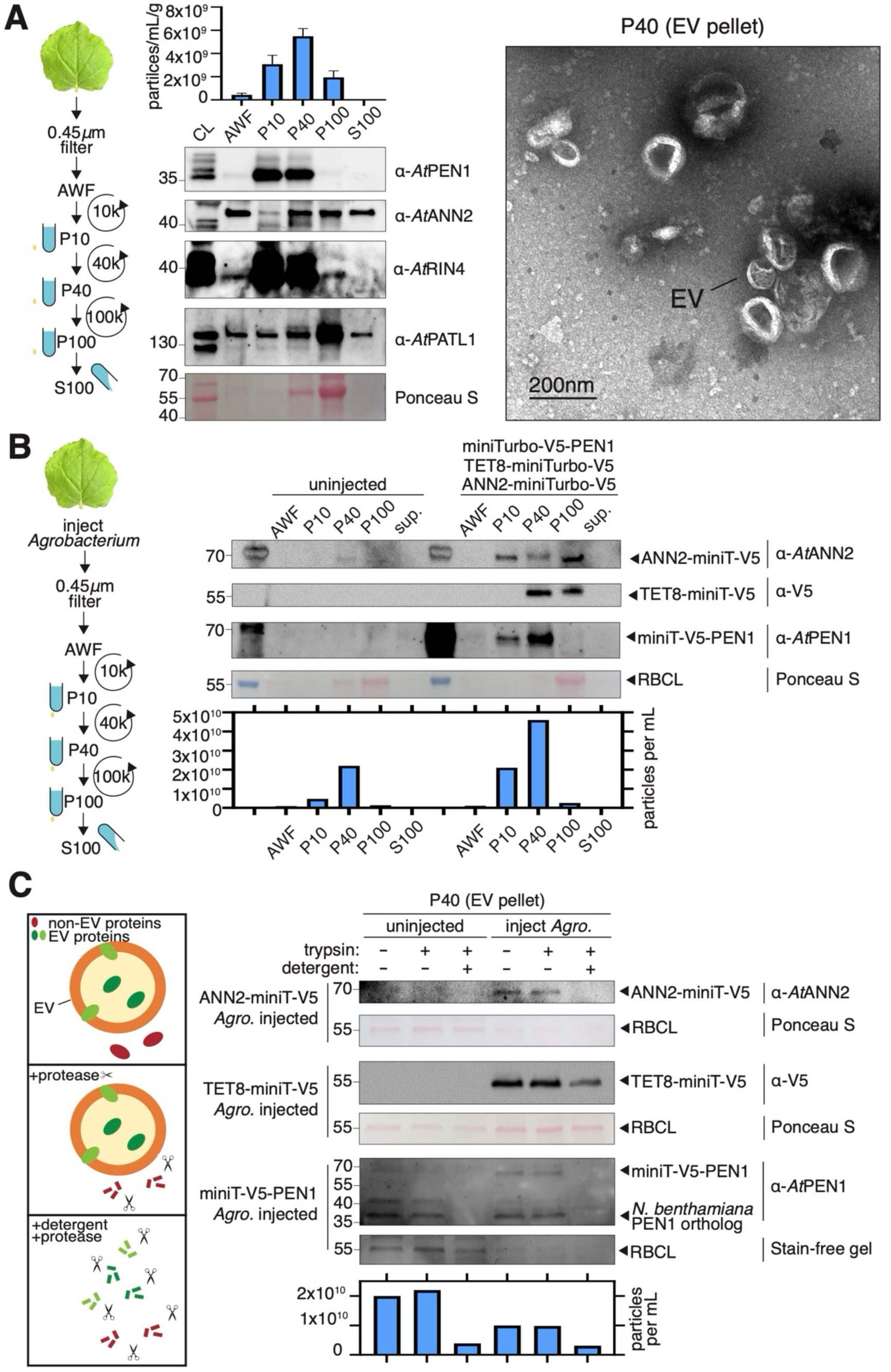
High speed pellets from *N. benthamiana* contain EVs. Nanoparticle tracking analysis (NTA), immunoblots, and negative-stain transmission electron micrographs (TEM) of various extracellular fractions from *N. benthamiana* leaves. **A)** Apoplastic wash fluid (AWF) was collected from the extracellular space of detached *N. benthamiana* leaves, filtered, and centrifuged at 10,000 × *g* to pellet large particles and cellular debris (P10). The supernatant of the P10 was centrifuged at 40,000 × *g* (P40) to pellet small particles. The supernatant of the P40 was centrifuged at 100,000 × *g* (P100). Error bars in NTA graph represent ±SEM from five independent biological replicates. TEM analysis of the P40 pellet revealed typical EV morphology with circular to ovoid shapes 50–200 nm in diameter and appearing concave due to dehydration of samples during negative staining. **B)** Extracellular fractions of uninjected *N. benthamiana* leaves, and leaves injected with a mixture of three different *Agrobacterium tumefaciens (Agro.)* strains carrying tagged versions of Arabidopsis EV marker proteins. Particles and transiently expressed Arabidopsis EV markers are identified in the P40, while Rubisco large subunit (RBCL) is more enriched in P100. **C)** Immunoblot and NTA following protease protection assay of EV pellet (P40). EV pellets were isolated from injected and uninjected *N. benthamiana* leaves. EV pellets were treated with detergent (1% TX-100 4°C 30 min) followed by protease (1µg/µL trypsin 37°C 1h), protease alone, or mock treatment. Digestion of EV marker proteins was assessed by immunoblot and disruption of EVs following detergent treatment was assessed by NTA. RBCL is partially resistant to trypsin digestion for unknown reasons. Fusion of the mini-Turbo tag onto EV marker proteins does not affect packaging into EVs.

### Identification of potential EV trafficking pathways using proximity labeling

To investigate potential biogenesis mechanisms for specific EV subpopulations, we used proximity labeling. We transiently expressed Arabidopsis EV marker proteins fused to the biotin ligase miniTurbo in *N. benthamiana*. To confirm that neither the miniTurbo ligase nor the V5 tag prevented the packaging of these fusion proteins into EVs, we isolated EVs and performed protease protection assays (Fig. 1B,C). Biotin ligase tagged versions of TET8, PEN1, and ANN2 all accumulated in the P40 pellet and were protected from digestion by trypsin in the absence of detergent, indicating that these Arabidopsis EV fusion proteins were secreted in *N. benthamiana* EVs.

To confirm that the EV-miniTurbo fusion proteins retained biotin ligase activity, we injected *N. benthamiana* leaves that were expressing each of the three different EV-miniTurbo fusion proteins individually with a biotin solution. After incubation, we isolated total soluble protein and detected biotinylated proteins using streptavidin linked to horse-radish peroxidase (strep-HRP) (Supplementary Fig. S2). We found that leaves expressing an EV-miniTurbo fusion protein had more biotinylated proteins than leaves that had only been injected with biotin solution, confirming the miniTurbo biotin ligase was active (Supplementary Fig. S2). Biotinylated proteins were also readily detected in EV pellets isolated from leaves expressing any of the miniTurbo fusion proteins, including the YFP-miniTurbo control, even though YFP-miniTurbo was not detected in these pellets (Supplementary Fig. S2). This latter observation suggests that proteins biotinylated in the cytoplasm can be loaded into EVs.

To identify candidate interacting proteins of TET8, PEN1, and ANN2, we first affinity-purified biotinylated proteins from whole cell lysates using streptavidin beads and confirmed enrichment of biotinylated proteins by probing with strep-HRP (Supplementary Fig. S2). We then used semi-quantitative mass spectrometry (MS) to analyze three independent biological replicates for TET8-, PEN1-, and ANN2-proximity labeled proteins isolated from cell lysates (Fig. 2, Supplementary Dataset S1). Candidate interacting proteins (enriched >2-fold relative to the YFP-miniTurbo and biotin only controls) of PEN1 and TET8 were enriched in gene ontology (GO) terms such as vesicle-mediated transport and vesicle tethering/docking/fusion. However, some GO terms were unique to PEN1 or TET8. TET8 candidate interacting proteins were uniquely enriched in endoplasmic reticulum (ER) processes and Golgi vesicle transport (Fig. 2).

**Supplementary Figure S2.**
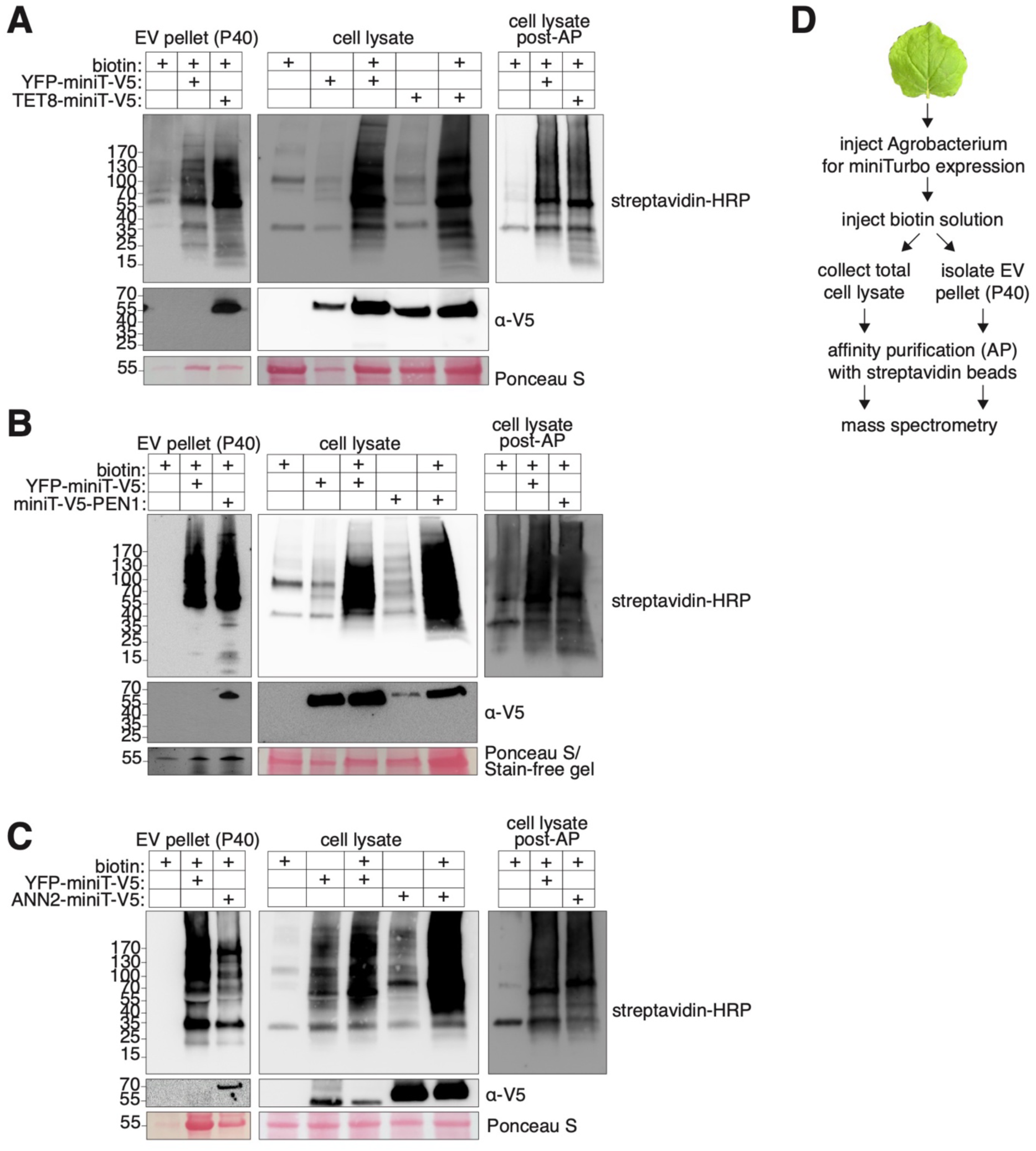
Proximity labeling by Arabidopsis EV marker proteins transiently expressed in *N. benthamiana.* **A)** TET8-miniTurbo. **B)** miniTurbo-PEN1. **C**) ANN2-miniTurbo. The fusion proteins were detected using anti-V5 antibody. Biotinylated proteins were detected using streptavidin-HRP (strep-HRP). **D)** Experimental work-flow for **A-C**. Biotin-labeled proteins were affinity-purified (AP) using streptavidin beads and analyzed by mass spectrometry (see Fig. 2 and Supplementary Fig. S3).

**Figure 2.**
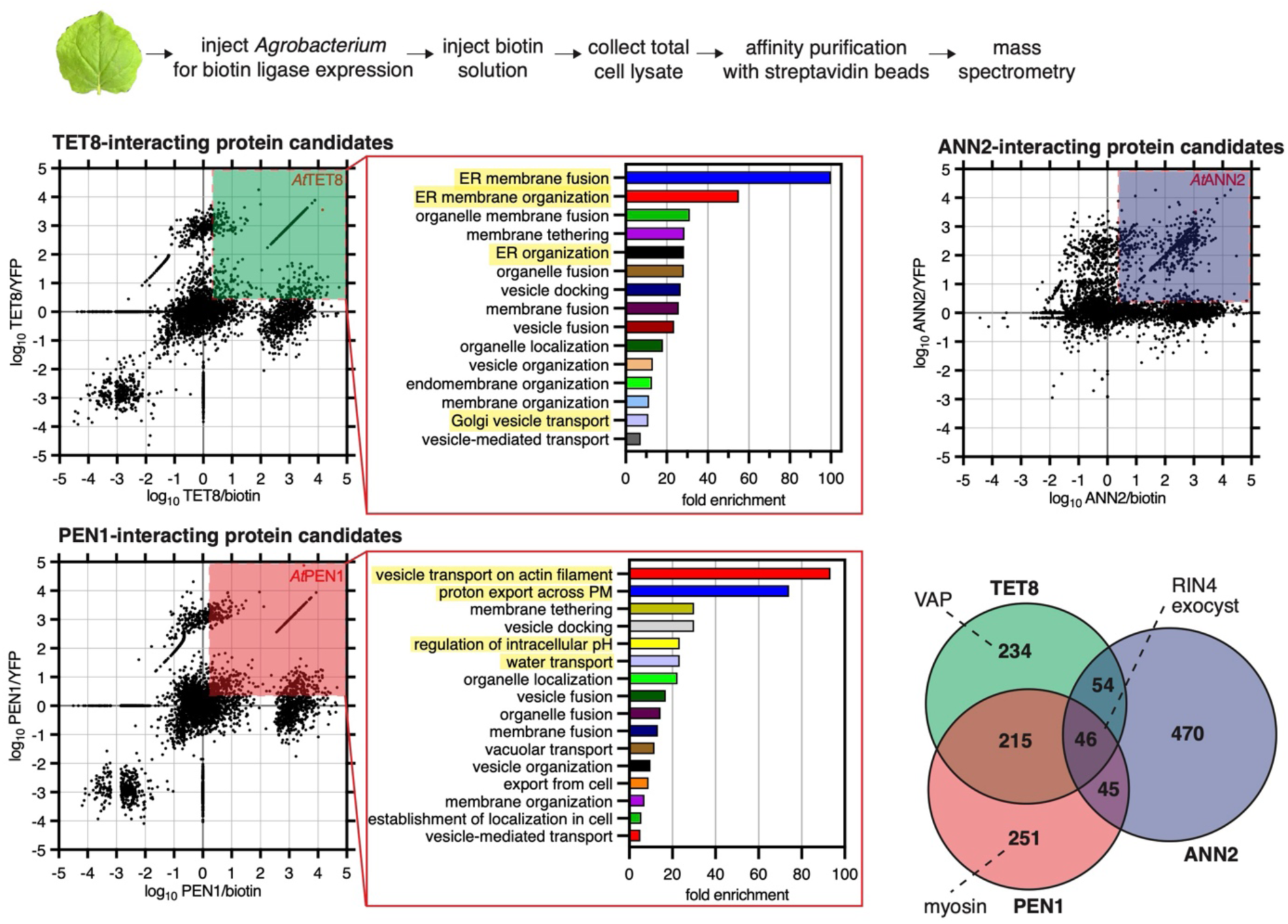
Proximity labeling in whole cell lysates implies distinct and shared proteins and pathways for EV subpopulation markers. Semi-quantitative mass spectrometry (MS) was performed on affinity-purified biotinylated proteins from *N. benthamiana* leaves transiently expressing Arabidopsis EV subpopulation marker proteins (see Supplementary Fig. 3). Enrichment plots show the intensity value of each protein identified by MS and its enrichment relative to both the YFP and biotin negative controls. Red boxes show candidate interacting proteins, defined as >2 fold enriched relative to both controls. Candidate interacting proteins were analyzed by GO and terms >5 fold enriched relative to expected were graphed. Highlighted GO terms were uniquely enriched for either PEN1- or TET8-candidate interacting proteins. The Venn diagram depicts the number and degree of overlap between the candidate interacting proteins for PEN1, TET8, and ANN2. The numbers indicate the amount of proteins that belong to each category. Only 46 proteins were common among all three EV protein candidate interactomes and included RIN4 and several exocyst proteins. Myosin proteins and VAP proteins were identified as PEN1- and TET8-candidate interacting proteins, respectively.

A previous study has described the interaction of TET8 with γ2-COPI at the Golgi (Liu et al., 2024). We found that several VAPs, which localize to the ER (Wang et al., 2016), uniquely interacted with TET8. In human cells, VAP proteins have been shown to drive the biogenesis of small, CD63+, RNA-containing EVs at MVB-ER contact sites (Barman et al., 2022; Barman et al., 2024). Since CD63 is structurally similar to TET8, TET8+ EVs biogenesis may also involve VAP proteins at MVB-ER contact sites (Boavida et al., 2013; Cai et al., 2018; Jimenez-Jimenez et al., 2019). On the other hand, PEN1 candidate interacting proteins were uniquely enriched in cytoskeleton and actin-related GO terms (Fig. 2). Although PEN1 and TET8 likely mediate endomembrane trafficking steps independent of EV biogenesis, this list of candidate interacting proteins should include those involved in EV biogenesis.

Previous studies have shown that PEN1 is recruited to actin patches in the plasma membrane during fungal infection, and that this PEN1 trafficking is disrupted when myosin genes are knocked out (Yang et al., 2014; Qin et al., 2021). We found that myosin proteins were also labeled by miniTurbo-PEN1 (Fig. 2). Actin and myosin have been linked to microvesicle secretion in the cilia of mammalian cells, but in plants it has been suggested that myosin transports endosomes to the plasma membrane (Kurth et al., 2017; Nager et al., 2017), raising the possibility that mysosin is involved in transport of MVBs to the plasma membrane, presumably by association with the limiting membrane of the MVB. Notably, PEN1 also interacted with proton and water transport-related proteins more often than TET8 (Fig. 2). This may be significant since these proteins are enriched in membrane nanodomains, suggesting that nanodomain organization of the plasma membrane may play a role in biogenesis of the PEN1+ EV subpopulation (Tapken and Murphy, 2015)(Fig. 2).

We also attempted to affinity-purify proteins from EV pellets, but only ANN2-miniTurbo EV pellets yielded informative results, likely due to very limited starting material for these analyses. We used gene ontology (GO) enrichment analysis and found that ANN2 interactors identified in EV pellets are likely to play a role in endosomal transport, including multivesicular bodies (MVBs), suggesting that ANN2 EVs may be derived from MVBs (Supplementary Fig. S3A). We found RIN4 and two exocyst complex proteins in ANN2-miniTurbo EV pellets, and these proteins were absent from both the “biotin only” and YFP-miniTurbo controls (Supplementary Fig. S5B). This result is consistent with our findings from the whole cell lysate proximity labeling, which also identified RIN4 and exocyst complex proteins as interacting with ANN2 (Fig. 2). The exocyst complex is an octameric tethering complex that is essential for conventional secretion from post-Golgi vesicles (Mei et al., 2018; Pereira et al., 2023).

When comparing the candidate interacting proteins of PEN1, TET8, and ANN2, there were surprisingly few candidate interactors shared by all three EV marker proteins (46 proteins out of 1,315 total candidate interacting proteins, <4%)(Fig. 2), likely reflecting that these proteins perform other cellular functions in addition to vesicle secretion. Candidate interactors shared by all EV subpopulation markers included RIN4, several exocyst component proteins, especially EXO70 family proteins, and some Rab GTPases.

In human cells, RNAi knockdown of EXO70 results in significantly lower EV secretion, and MVBs build up in the cell, unable to fuse with the plasma membrane (Liu et al., 2023). RIN4 has no known human homologs. RIN4 has been shown previously to interact with EXO70 proteins (Mackey et al., 2003). Together, these results may suggest that 1) MVBs are the main source of EVs in plants, 2) the exocyst complex, RIN4, and Rab proteins play a central role in EV secretion, and 3) that TET8 EV biogenesis involves VAP proteins while PEN1 EV biogenesis involves myosin proteins.

**Supplementary Figure S3.**
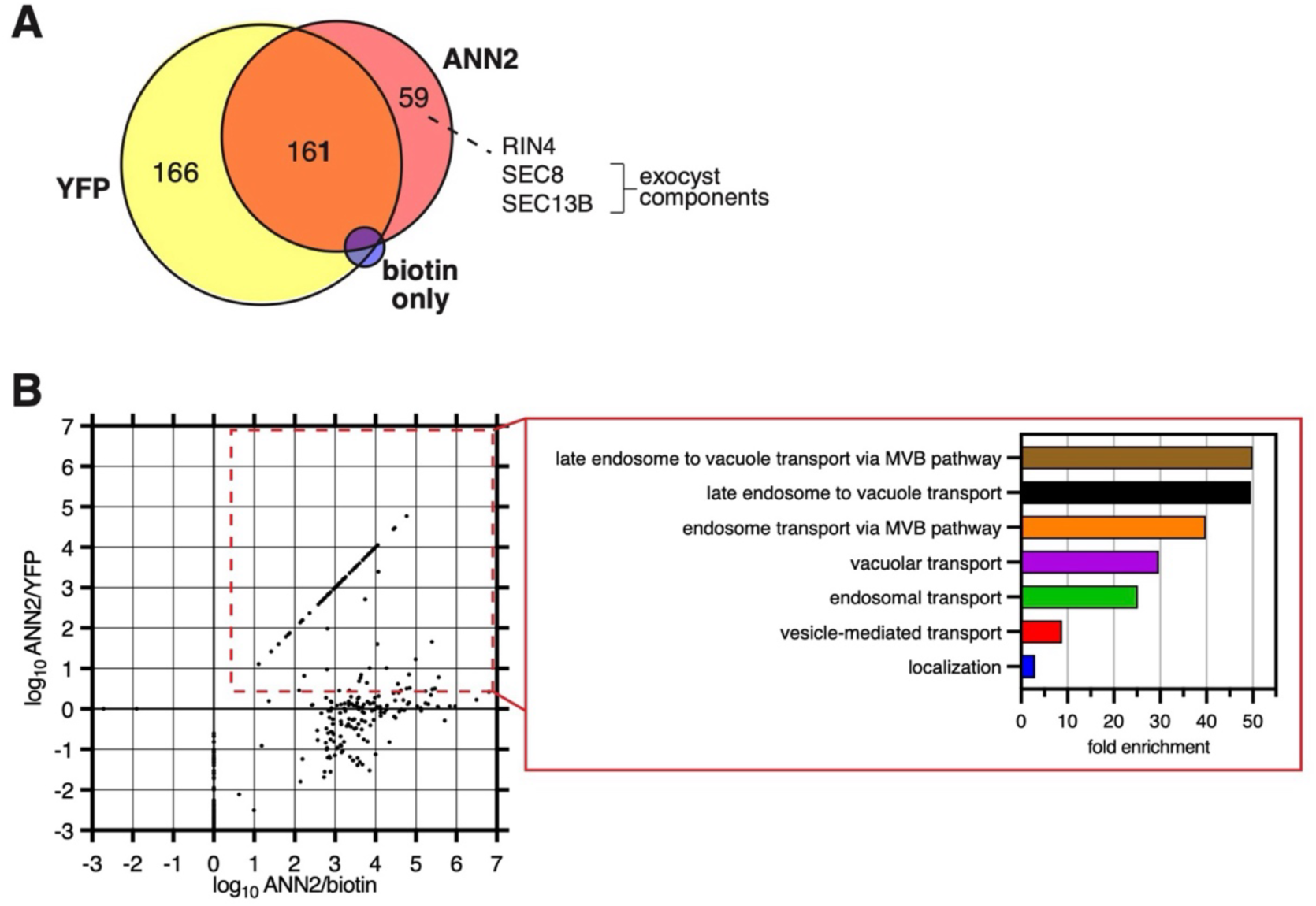
Proteins secreted in EVs following proximity labeling of ANNEXIN2 (ANN2) suggests secretion of ANN2+ EVs involves multivesicular bodies (MVBs). **A)** Semi-quantitative mass spectrometry (MS) was performed on affinity-purified biotinylated proteins from the EV pellet obtained from *N. benthamiana* leaves transiently expressing ANN2, a YFP control, or no construct/biotin only injection (see Supplementary Fig. S2)). Enrichment plots show the intensity value of each protein identified by MS and its enrichment relative to both the YFP and biotin negative controls. Red boxes show candidate interacting proteins, defined as >2 fold enriched relative to both controls. Candidate interacting proteins were analyzed by GO. **B)** Venn diagram of biotinylated proteins identified in each EV pellet dataset. Proteins uniquely identified in the ANN2 sample included RIN4 and exocyst proteins. Proteins identified in the YFP sample were likely biotinylated before secretion. Very few proteins were identified in the biotin only control sample, which is consistent with the blots in Supplementary Fig. S2.

### The exocyst complex interacts and co-localizes with EV subpopulation markers including RIN4 on MVB-like structures

To further investigate the role of EXO70 proteins and the exocyst complex in the secretion of EVs, we transiently co-expressed EXO70B2 with TET8, PEN1, or ANN2 in *N. benthamiana* leaves and assessed potential interactions using co-immunoprecipitation (co-IP). We found that while PEN1 and TET8 strongly interacted with EXO70B2 in *N. benthamiana* co-IPs, ANN2 did not interact with EXO70B2 significantly above that observed in the negative control (Fig. 3A). Consistent with this, we observed co-localization of TET8 with EXO70B2 in cytoplasmic puncta, suggesting that EXO70B2 may interact with TET8+ in the cytoplasm prior to tethering to the plasma membrane and secretion of TET8+ EVs (Fig. 3B). Together, these results suggest that EXO70 proteins may contribute to the secretion of TET8+ and PEN1+ EVs, but their role in secretion of ANN2+ EVs may be more indirect.

**Figure 3.**
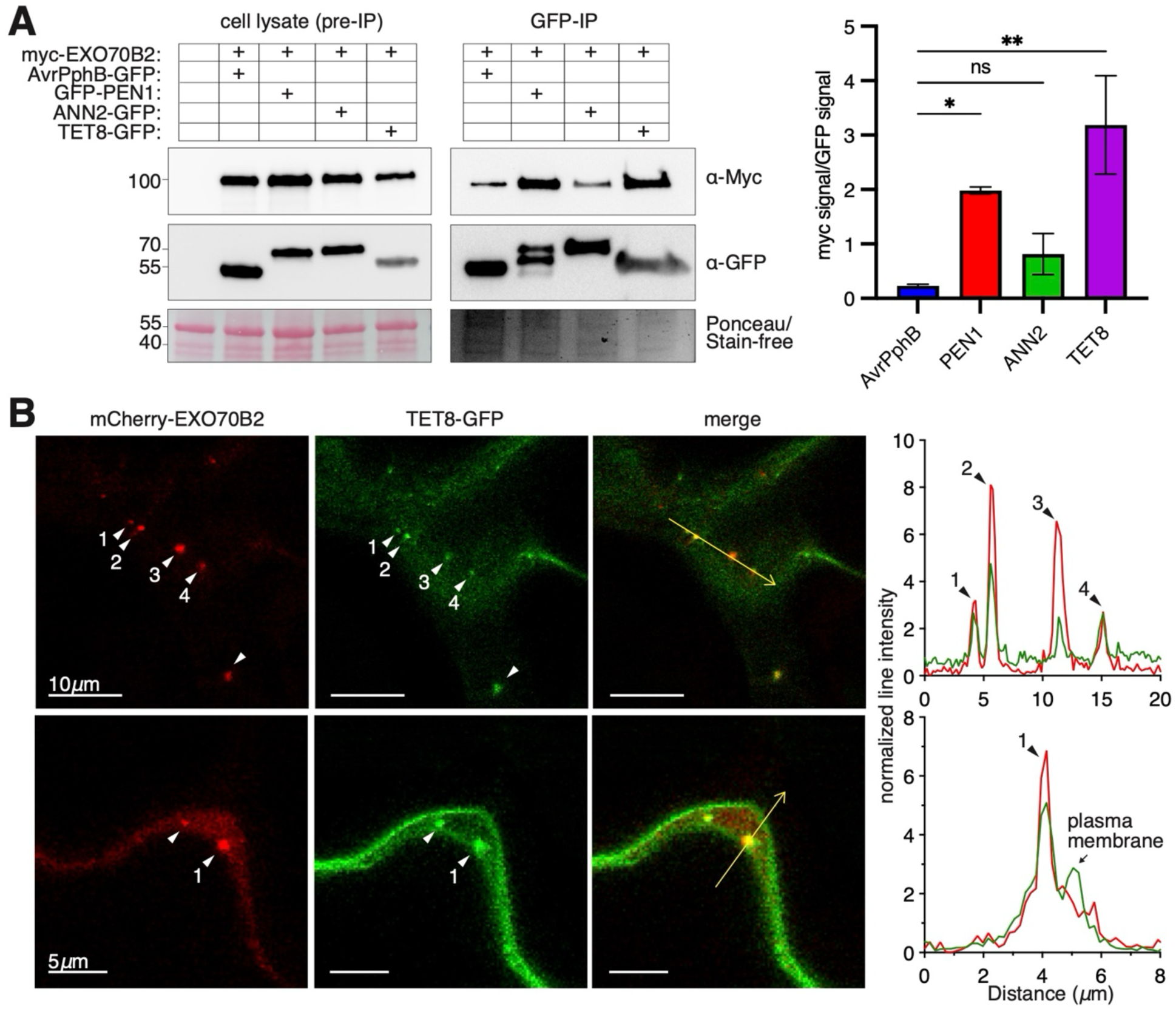
Interaction of EXO70B2 with EV subpopulation marker proteins. **A**) Co-immunoprecipitation (co-IP) of EXO70B2 with EV marker proteins using transient expression in *N. benthamiana*. AvrPphB-GFP is a plasma-membrane targeted protein that was used as a negative control. The amount of GFP and myc signal was quantified using densitometry, and the ratio of myc to GFP signal was graphed as an average of three independent biological replicates. The error bar indicates ± standard deviation, and statistical significance was detected using a one-way ANOVA followed by a post-hoc Tukey’s test, with p-value of <0.05 shown by *; <0.01 by **; ns is not statistically significant. **B**) Confocal fluorescence microscopy of proteins transiently expressed in *N. benthamiana*. Top and bottom rows show two different lobes of a single pavement cell. White arrowheads indicate instances of co-localization. Lines for line intensity plots are indicated in images by yellow vectors. Normalized intensity of GFP and mCherry signal was plotted along each vector, showing multiple instances of co-localization, as indicated.

We proceeded to test the hypothesis that RIN4 also has a general role in EV secretion. We observed that RIN4 localized to the plasma membrane but was also found in large (median diameter = 1.1μm), circular, cytoplasmic structures (Fig. 4A). These structures sometimes appeared to fuse with the plasma membrane. RIN4 seemed to preferentially associate with the limiting membrane of these structures, suggesting that RIN4 is present at the surface of these structures. These RIN4 structures also co-localized with EXO70B2 (Fig. 4B). RIN4 also co-localized with EXO70B2 more often than other EV marker proteins (Supplemental Fig. S4). The physical interaction between RIN4 and EXO70 family proteins has previously been shown using yeast two hybrid assay (Redditt et al., 2019), and this interaction between RIN4 and EXO70 proteins is supported by our microscopy observations. Together, these results suggest that RIN4 localizes with EXO70B2 to the surface of large circular organelles adjacent to the plasma membrane, which may represent MVBs or paramural bodies (MVBs that have fused with the plasma membrane). However, we were unable to detect either spatial interaction or physical association of RIN4 with TET8, PEN1, or ANN2 (beyond co-localization of RIN4, TET8, and PEN1 in the plasma membrane) suggesting that the nature of the interaction between RIN4 and other EV marker proteins is transitory.

**Figure 4.**
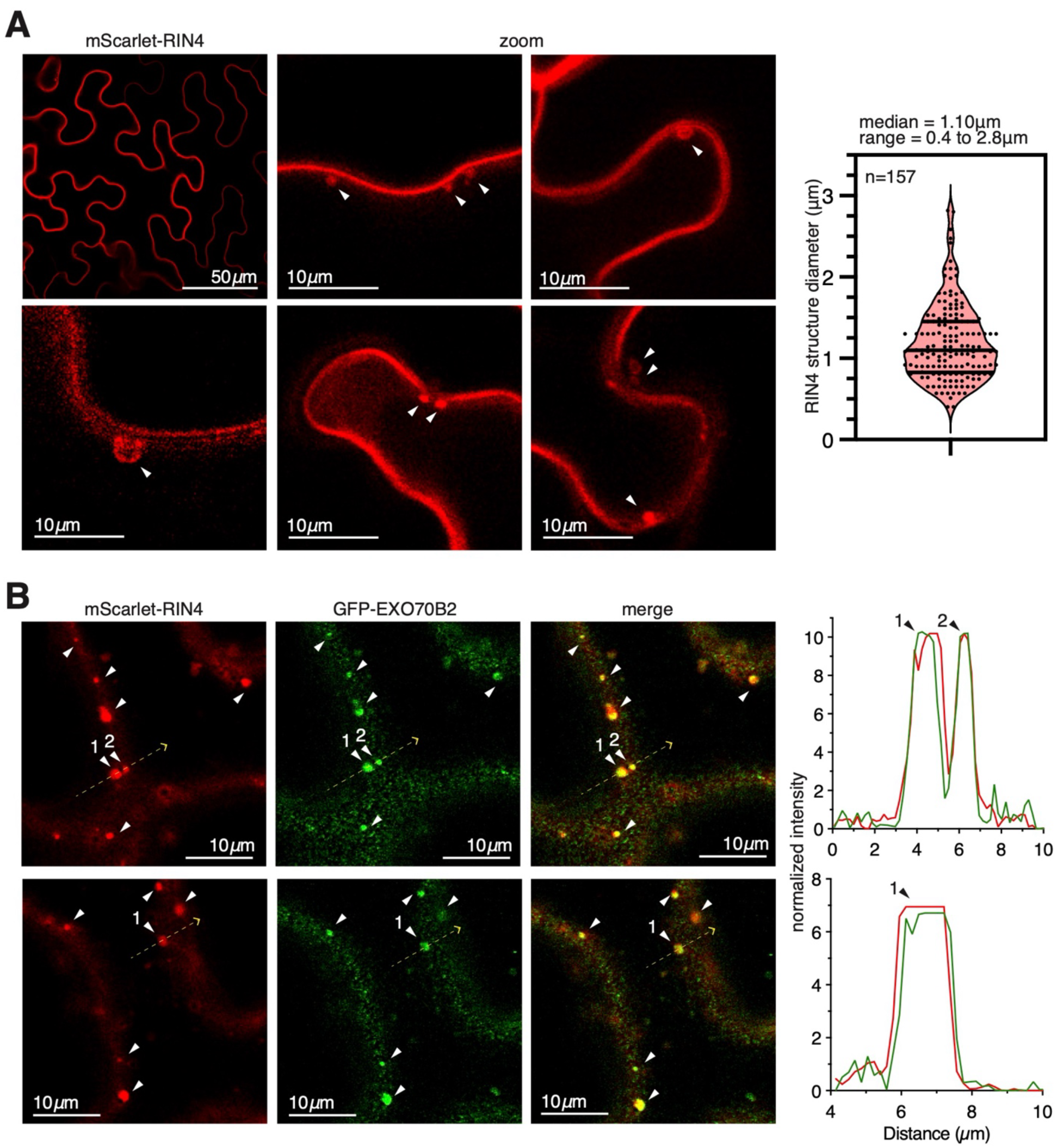
RIN4 and EXO70B2 interact on large, circular, cytoplasmic structures. **A**) Confocal fluorescence microscopy of *N. benthamiana* leaves transiently expressing mScarlet-RIN4. Arrowheads indicate large, circular, sometimes hollow, cytoplasmic structures. The diameter of these structures was measured using Fiji and graphed as a violin plot, with dark lines representing the median and the upper- and lower-quartile range. **B**) Confocal fluorescence microscopy of *N. benthamiana* leaves transiently expressing both mScarlet-RIN4 and GFP-EXO70B2. Arrowheads indicate co-localization of the two proteins in large, cytoplasmic puncta. Intensity of signal for mScarlet-RIN4 and GFP-EXO70B2 was analyzed along each white dotted line and plotted in red and green respectively.

**Supplementary Figure S4.**
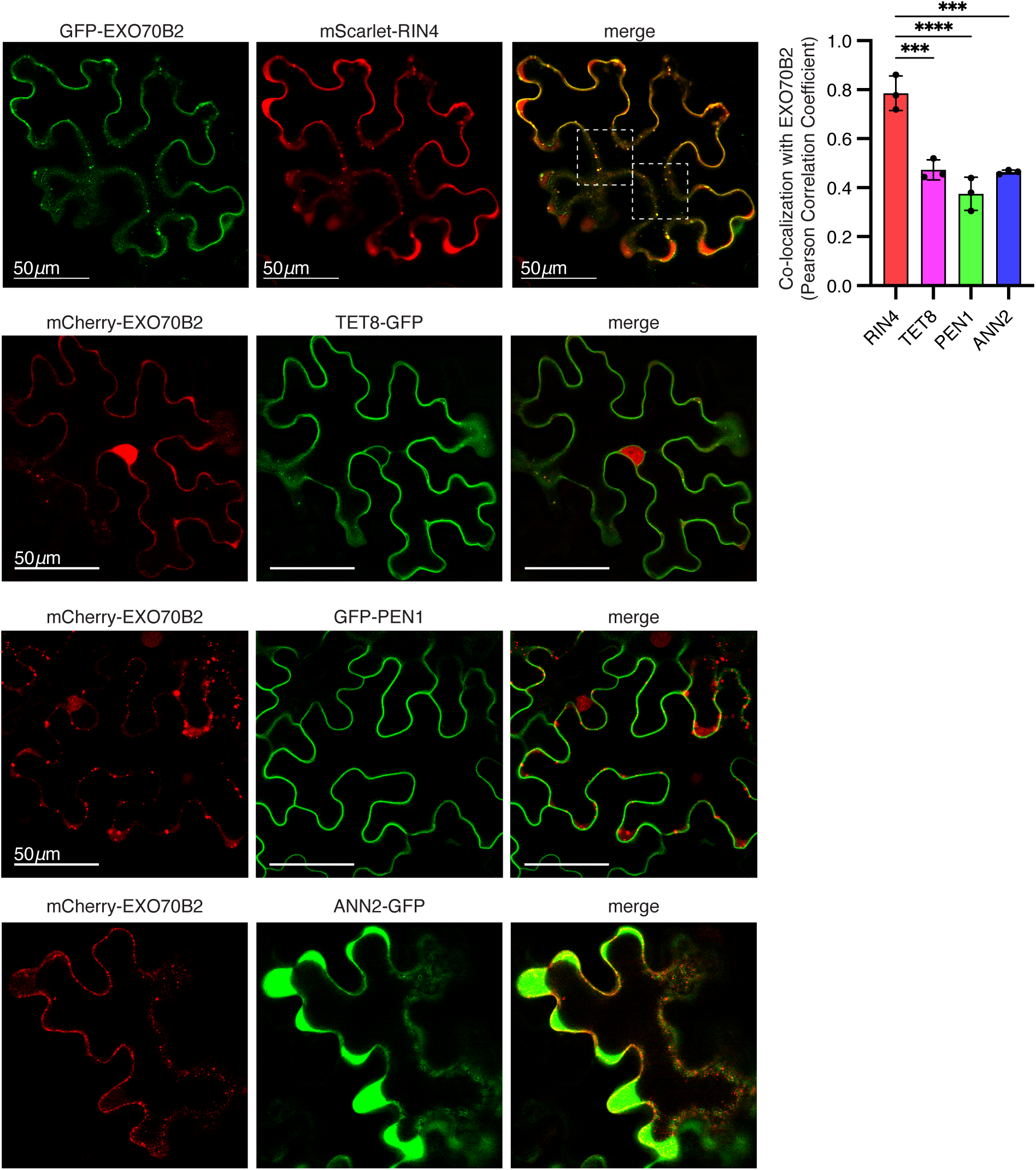
EXO70 co-localizes with RIN4 to a greater degree than other EV marker proteins. Confocal fluorescence microscopy of *N. benthamiana* leaves transiently expressing fluorescent tagged version of EXO70B2 and the EV marker proteins RIN4, TET8, PEN1, and ANN2. White dotted-line boxes are enlarged in Figure 3. Pearson correlation co-efficient of the amount of overlap between EXO70B2 and various EV marker proteins was calculated for three cells. Error bars indicate ± standard deviation. Statistical significance was detected using a one-way ANOVA followed by a post-hoc Tukey’s test, with p-value of <0.01 is shown by *** and <0.001 by ****.

### PEN1 and TET8 localize to overlapping populations of MVB-like structures

We expressed TET8 in *N. benthamiana* and found that, as previously reported, TET8 localized to the periphery and lumen of MVB-like structures (Fig. 5A) (He et al., 2021; Liu et al., 2024). We found that PEN1 localized to similarly-sized large structures (median diameter = 1µm), both when transiently expressed in *N. benthamiana* with a strong promoter and when stably expressed in Arabidopsis with a native promoter (Fig. 5B and C). These structures were reminiscent of the large, vesicle-like structures that have been observed in association with fungal papillae (Zeyen and Bushnel, 1979; Assaad et al., 2004; An et al., 2006), where PEN1 has previously been shown to localize (Collins et al., 2003; Nielsen et al., 2012). We reasoned that since TET8 and PEN1 are secreted in different subpopulations of EVs, that they would likely be secreted from distinct classes of MVBs. However, when we observed MVB-like structures in a stable transgenic Arabidopsis line expressing RFP-PEN1 and TET8-GFP, we found that PEN1 and TET8 co-localized to the same MVB-like structures (Fig. 5D, Supplementary Fig. S5). This result suggests that both TET8 and PEN1 EVs may be formed on the same type of MVBs, and that the secretion of TET8 and PEN1 in separate subpopulations may instead depend on differing mechanisms of ILV formation.

**Figure 5.**
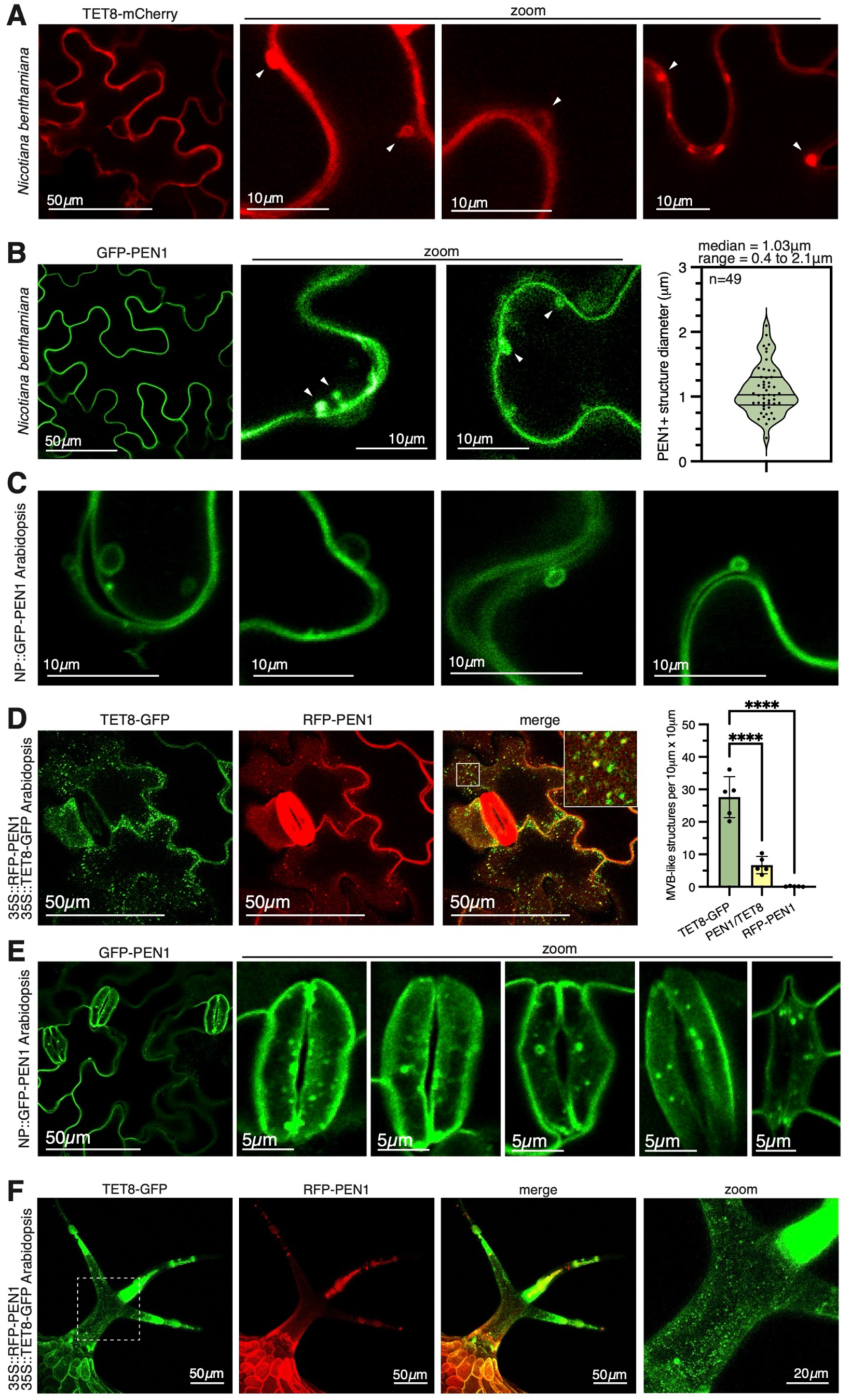
TET8 and PEN1 localize to MVB-like structures. **A)** Transient expression of TET8-mCherry in *N. benthamiana* leaves. White arrowheads indicate MVB-like structures. **B)** Transient expression of GFP-PEN1 in *N. benthamiana leaves.* The diameter of MVB-like structures was measured using Fiji and graphed as a violin plot, with dark lines representing the median and the upper- and lower-quartile range. **C**) Confocal fluorescence microscopy of Arabidopsis pavement cells stably expressing native PEN1 promoter (NP) fused to GFP-PEN1. Note large circular structures, similar to those observed in *N. benthamiana,* were most frequently observed in older leaves **(D)** Confocal fluorescence microscopy of epidermal leaf cells of a stable transgenic Arabidopsis line expressing TET8-GFP and RFP-PEN1, each with 35S promoters. Localization of GFP and RFP signal to MVB-like structures was manually counted for five independent images, and co-localization events were quantified using Fiji. Statistical significance was detected with a one-way ANOVA, followed by Tukey’s post-hoc test, with a p-value of <0.0001 shown by ****. Together, this shows that TET8 localizes MVB-like structures about 5 times more often than PEN1, and when PEN1 appears in MVB-like structures, it nearly always co-localizes with TET8. Note also the localization of PEN1 to the stomata, and the absence of TET8 in the stomata. Additional images included in Supplemental Fig. S5. The inset image is outlined by a white box and is 5 microns x 5 microns. **(E)** Confocal fluorescence microscopy of Arabidopsis stomata stably expressing GFP-PEN1 under its native promoter. GFP signal in stomata was detected in large MVB-like structures similar to that in leaf epidermal cells in **(B)** and **(C)**. MVB-like structures were especially prevalent near the stomatal pore or aperture. **(F)** Confocal fluorescence microscopy of trichomes or leaf hairs in Arabidopsis stably expressing TET8-GFP and RFP-PEN1. TET8-GFP puncta are observed throughout the central trichome stalk. GFP signal is remarkably strong in the trichome branches, where some RFP signal is also observed.

We observed that PEN1 is strongly expressed in guard cells, and that PEN1 MVB-like structures are apparent in guard cells near the stomatal aperture (Fig. 5E). This suggests that guard cells may be a source of PEN1 EVs. We also observed that TET8 endosomes were present in trichomes or leaf hairs, suggesting that trichomes may be a source of TET8 EVs (Fig. 5F). These results suggest that specific tissues may differ in their secretion of EV subpopulations.

Gene ontology (GO) analysis of TET8 and PEN1 candidate interactors indicated that in addition to the enrichment of secretion-related GO terms, TET8 interactors included endoplasmic reticulum (ER)-related GO terms while PEN1 candidate interactors included cytoskeleton-related GO terms (Fig. 2). We proceeded to test the involvement of these specific mechanisms in the biogenesis of TET8+ and PEN1+ EVs, respectively. We found that co-expression of VAP27 with TET8 altered the localization of TET8, causing it to be retained in the ER in numerous medium-sized punctate structures that also included VAP27 (Supplementary Fig. S6A). In contrast, PEN1 did not appear to be retained in the ER upon VAP27 expression, and VAP27 co-localized with TET8 significantly more frequently than PEN1 (Supplementary Fig. S6A). This suggests that, like CD63+ EVs in human cells, TET8+ EV biogenesis might originate from MVB-ER contact sites, while PEN1+ EV biogenesis does not involve this mechanism (Barman et al., 2022). Instead, we found that PEN1 interacted with the globular tail domain (GTD) of Myosin XI-K, consistent with the enrichment of cytoskeleton-related proteins in the PEN1 sample of the proximity labeling experiment (Supplementary Fig. S6B, Fig. 2). These results suggest that while TET8 and PEN1 co-localize to the same MVB-like structures, some aspects of their biogenesis remain distinct.

**Supplementary Figure S5.**
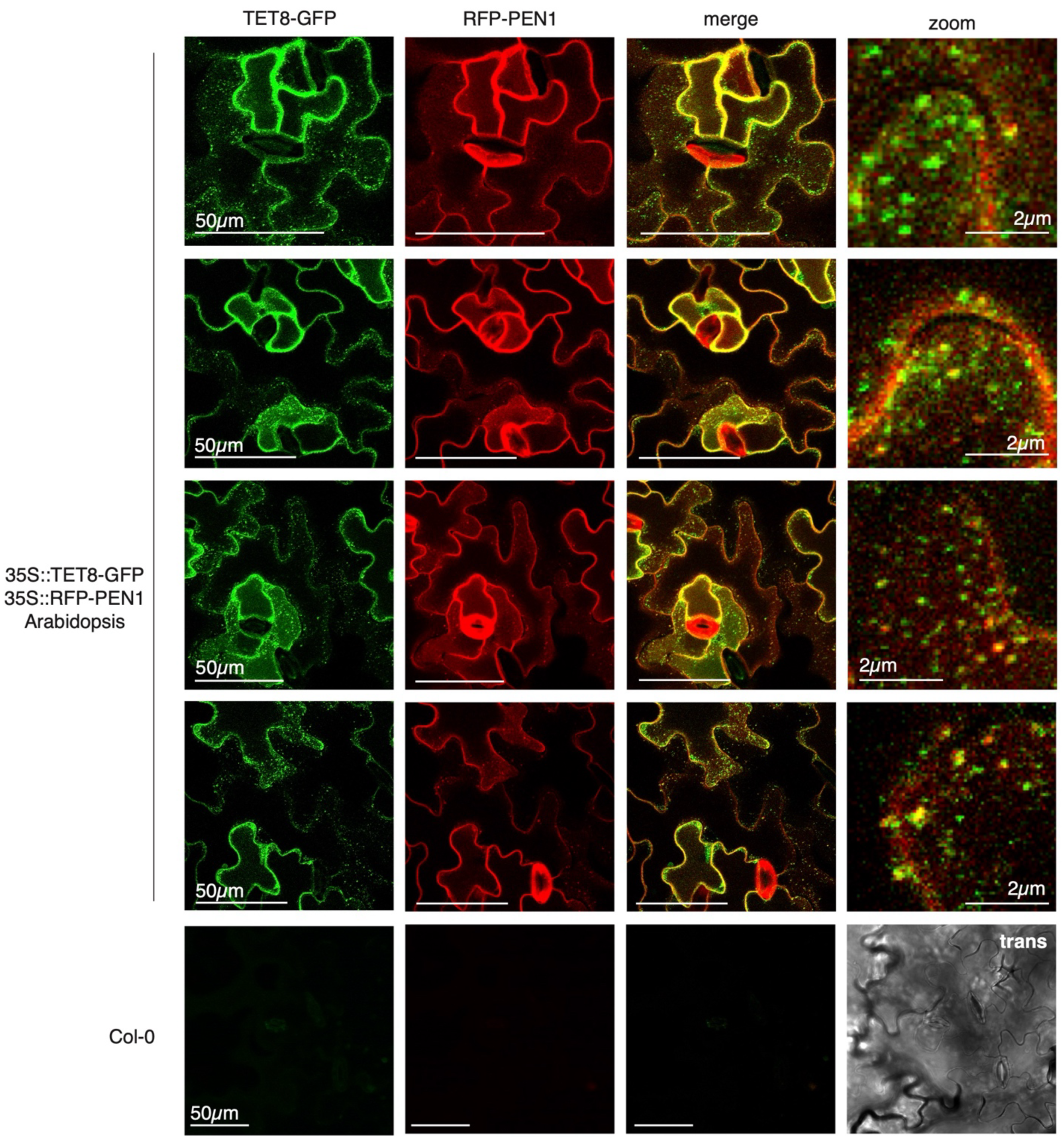
TET8 and PEN1 co-localize to MVB-like structures. Confocal fluorescence microscopy images of leaf epidermal cells from 4-week old Arabidopsis seedlings stably expressing TET8-GFP and RFP-PEN1 transgenes. Wild-type seedling with the same microscope settings and image processing are shown as a control. These representative images and others were used for co-localization analysis graphed in Fig. 5D.

**Supplementary Figure S6.**
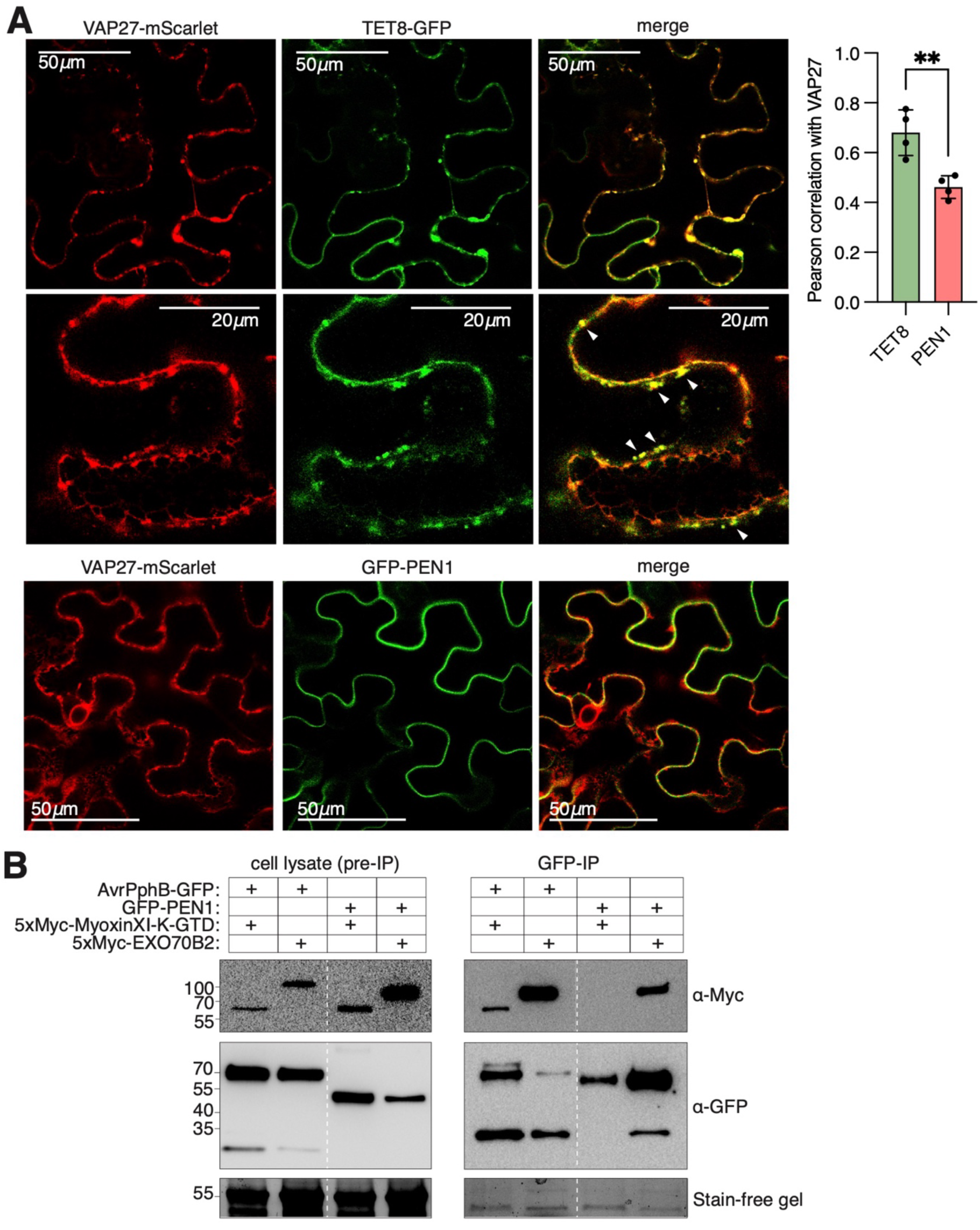
Biogenesis of TET8+ and PEN+ EVs involves VAP and myosin proteins respectively. **A)** VAP27 and TET8 co-localize when co-expressed in *N. benthamiana.* White arrowheads indicate colocalization in puncta. **B)** TET8 partially localizes to the ER. To quantify colocalization of TET8 with an ER marker, with VAP27, and with PEN1, a Pearson correlation coefficient was calculated for 4-5 cells per expression combination and statistical significance was detected using a one-way ANOVA and a Tukey post-hoc test, with p-value of <0.05 shown by *; <0.01 by **; and <0.0001 by ****.

### Autophagy is indirectly involved in the secretion of TET8+ EVs

Previous reports in human cell culture have shown that amphisomes mediate secretory autophagy (Jeppesen et al., 2019; Leidal et al., 2020), and a recent report in plants showed that amphisomes form in plants prior to fusion with the vacuole for degradation (Jeppesen et al., 2019; Leidal et al., 2020; Zhao et al., 2022). We investigated whether amphisomes could also form prior to secretion of EVs. We found that the small GTPase ARA6 co-localized with the autophagosome marker ATG8a in *N. benthamiana* (Fig. 6A). ARA6 is a marker of secretory MVBs and has been shown to participate in defense-related secretory membrane trafficking, localizing to sites of fungal penetration (Ebine et al., 2011; Nielsen et al., 2012), which suggests that at least some ARA6 MVBs are destined for secretion. Given our finding that ARA6 co-localizes with ATG8a, it seems plausible that amphisomes could form prior to secretion.

**Figure 6.**
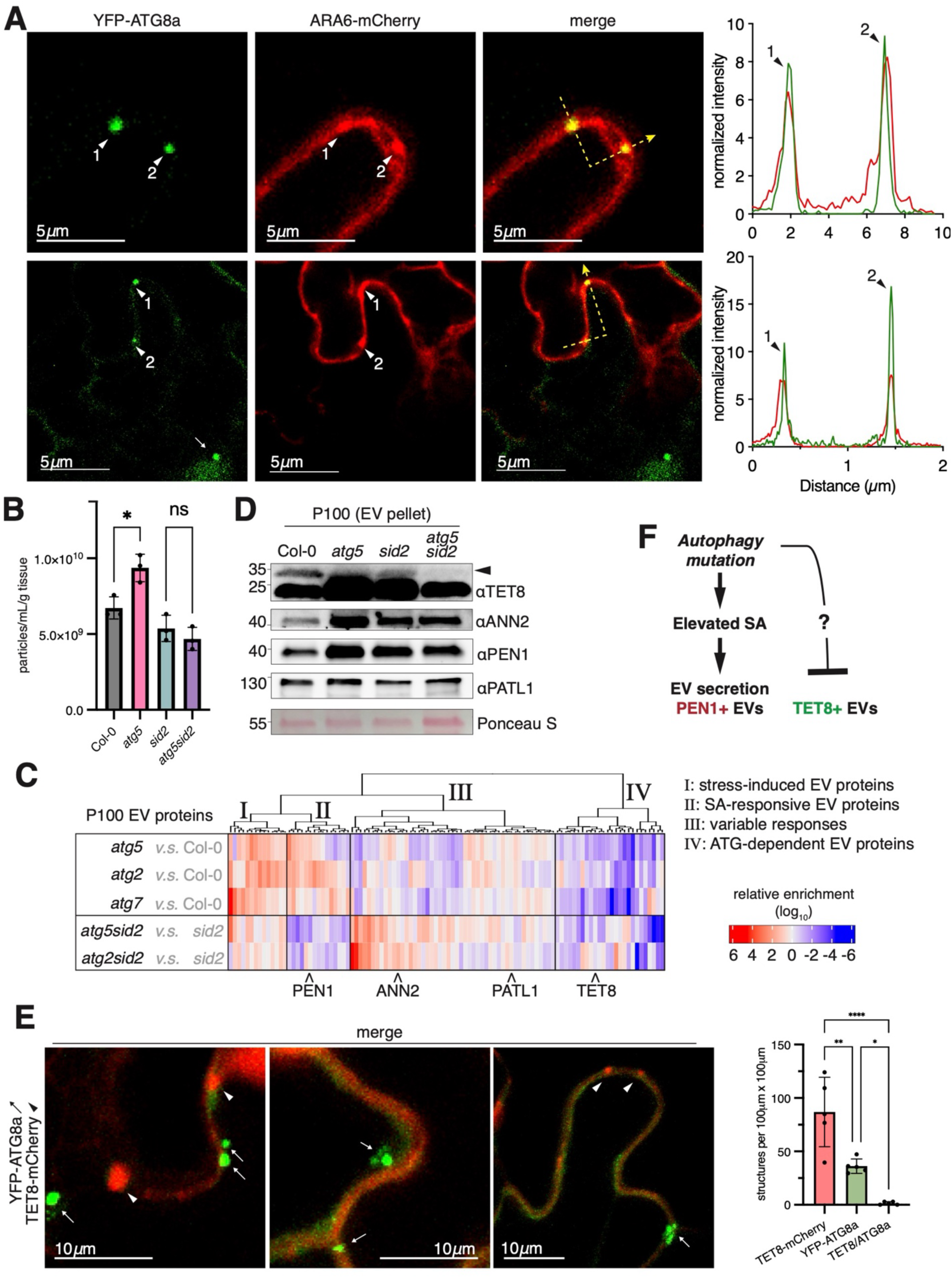
Autophagy indirectly influences the secretion of TET8+ EVs. **A)** YFP-ATG8 and ARA6 mCherry partially co-localize in puncta when transiently expressed in *N. benthamiana*. The dotted yellow lines indicate the lines used for the intensity plot. The white arrowheads indicate points of co-localization, while the white arrow indicates a point of YFP signal that lacks mCherry signal. **B)** Nanoparticle tracking analysis (NTA) of EV pellets (P100) isolated from various genotypes. Three independent biological replicates are graphed, and statistical significance was detected by a one-way ANOVA followed by Tukey post-hoc test, p-value of <0.05 is shown by *; ns is not significant. **C)** Semi-quantitative mass spectrometry (MS) of EV proteins (P100) of various genotypes is visualized as a heatmap. Only proteins that were present in a high-quality EV proteome are shown (Koch et al., 2025). The enrichment values were organized by hierarchical clustering into four general clades, as indicated by roman numerals. Clade I represents proteins that are increased in the EV pellet when the autophagy pathway is mutated, possibly due to cellular stress. Clade II proteins were increased in the EV pellet when autophagy was mutant but returned to normal levels when salicylic acid levels were reduced, suggesting that these EV proteins are secreted in response to high levels of salicylic acid. In accordance with this, PEN1 which has been shown to be secreted in EVs in response to salicylic acid, belongs to Clade II. Clade III proteins are generally enriched or unchanged in *atg sid2* double mutants compared to *sid2* single mutants, thus are not dependent on autophagy for secretion. Clade IV proteins are decreased in the EV pellet when the autophagy pathway is mutant, and thus potentially depend on autophagy for secretion. **D)** Immunoblot of EV pellet (P100) from various mutant genotypes. The arrowhead indicates the TET8-specific band at ∼35 kDa (a strong cross-reacting band is present at ∼25 kDa). Note depletion of TET8 in EV pellet of the *atg5 sid2* mutant compared to the *sid2* mutant alone. **E)** TET8-mCherry does not appear to localize to autophagosomes. TET8-mCherry was transiently co-expressed with the autophagosome marker YFP-ATG8a in *N. benthamiana.* Leaves were placed in the dark for 2 days prior to imaging to induce autophagosome formation. Images are maximum projections of 2 µm z-stacks to help visualize ATG8a signal. ATG8a puncta (indicated by arrows) likely represent autophagosomes while TET8 puncta (indicated by arrowheads) likely represent TET8+ MVBs. Subcellular structures were manually identified in five independent images, and their co-localization was determined with Fiji. Statistical significance was detected using a one-way ANOVA followed by a Tukey’s post-hoc test, with p-value of <0.05 shown by *; <0.01 by **; and <0.0001 by ****. **F)** Model for impact of autophagy mutations on secretion of EV subpopulations. PEN1+ EVs are increased in autophagy mutants due to increase in SA, since a *sid2* mutant background abolished this increase. TET8+ EVs are decreased in autophagy mutants, but there is no evidence of co-localization of TET8+ MVBs with autophagosomes, suggesting an unknown pleiotropic effect is responsible for the depletion of TET8+ EVs in autophagy mutants.

To investigate the contribution of autophagy and amphisomes to secretion of EVs in healthy plants, we collected an EV pellet from wild-type and several autophagy deficient mutants, *atg5, atg7, and atg2.* When comparing the number of particles in EV pellets, we found that contrary to our hypothesis, autophagy mutations resulted in significantly increased particle secretion (Fig. 6B). Previous studies have shown that autophagy mutations, especially in mature plants, have many pleiotropic effects, including the accumulation of salicylic acid (SA) (Yoshimoto et al., 2009). Foliar spray of SA has been shown to increase secretion of EVs and some EV subpopulations (Rutter and Innes, 2017). Therefore, we used a *sid2* mutant background to avoid accumulation of SA (Nawrath and Metraux, 1999; Yoshimoto et al., 2009). Indeed, we found that autophagy mutants in the *sid2* background did not secrete excess particles, suggesting that the increase in particles in autophagy mutants is at least partially due to a pleiotropic effect (Fig. 6B).

To assess which EV subpopulations, if any, were affected by autophagy mutations, we analyzed EVs isolated from autophagy mutants in both wild-type and *sid2* mutant backgrounds using mass spectrometry and found that only a small group of EV proteins were depleted from autophagy mutants regardless of background (Fig. 6C and Supplementary Dataset S2). Interestingly, this group of autophagy-dependent EV proteins included TET8, a finding which we confirmed by immunoblotting (Fig. 6D). To investigate whether autophagy was directly involved in the secretion of TET8+ EVs through formation of amphisomes, we observed the localization of TET8 and ATG8a in *N. benthamiana* (Fig. 6E). We could find no evidence of their co-localization in healthy plants, suggesting that autophagy indirectly contributes to the secretion of TET8+ EVs through an unknown mechanism (Fig. 6F).

### EV secretion mutants are more susceptible to infection by *C. higginsianum*

To identify Arabidopsis mutations that impair EV secretion, we used a reverse genetic screen directed by our proximity labeling study. We selected exocyst mutants particularly in the EXO70 family and *rin4*, as these proteins were top candidates from the proximity labeling experiment and follow-up experiments. We also made use of dominant negative and conditional alleles to test the effects of Rab GTPase knockdown on EV secretion, as Rab GTPases were commonly identified by proximity labeling and have previously been shown to be secreted with Arabidopsis EVs (Fig. 2) (Rutter and Innes, 2017; Koch et al., 2025). We also selected mutants specific for EV subpopulations, such as *vap27* and *tet8* mutants. Where possible, we selected mutants which had little-to-no pleiotropic phenotypes because EV secretion is increased during stress. We also selected lines that were mutant for multiple members of the same family because similar proteins often have redundant functions.

Among the Rab GTPases identified in the EV proteome, we first focused on RabA family member RabA2a (At1g09630) as this Rab has previously been shown to interact with PEN1/SYP121 and to regulate formation of a SNARE ternary complex at the plasma membrane (Pang et al., 2022). Because there are multiple *RabA* family members in Arabidopsis that appear to be at least partially redundant in function, we made a dominant negative allele of *RabA2a* (*RabA2a(N125I*)) under control of an estradiol inducible promoter. We found that PEN1+ EV secretion was strongly suppressed when RabA family function was disrupted (Fig. 7A, Supplementary Fig. S7A). Even without spraying estradiol, fewer PEN1+ EVs were secreted, suggesting that this construct expresses at least some RabA2a in the absence of estradiol. In sharp contrast, TET8+ EVs were increased when the RabA family was disrupted (Fig. 7A). This result is consistent with the model that PEN1 and TET8 are secreted through distinct cellular processes.

**Figure 7.**
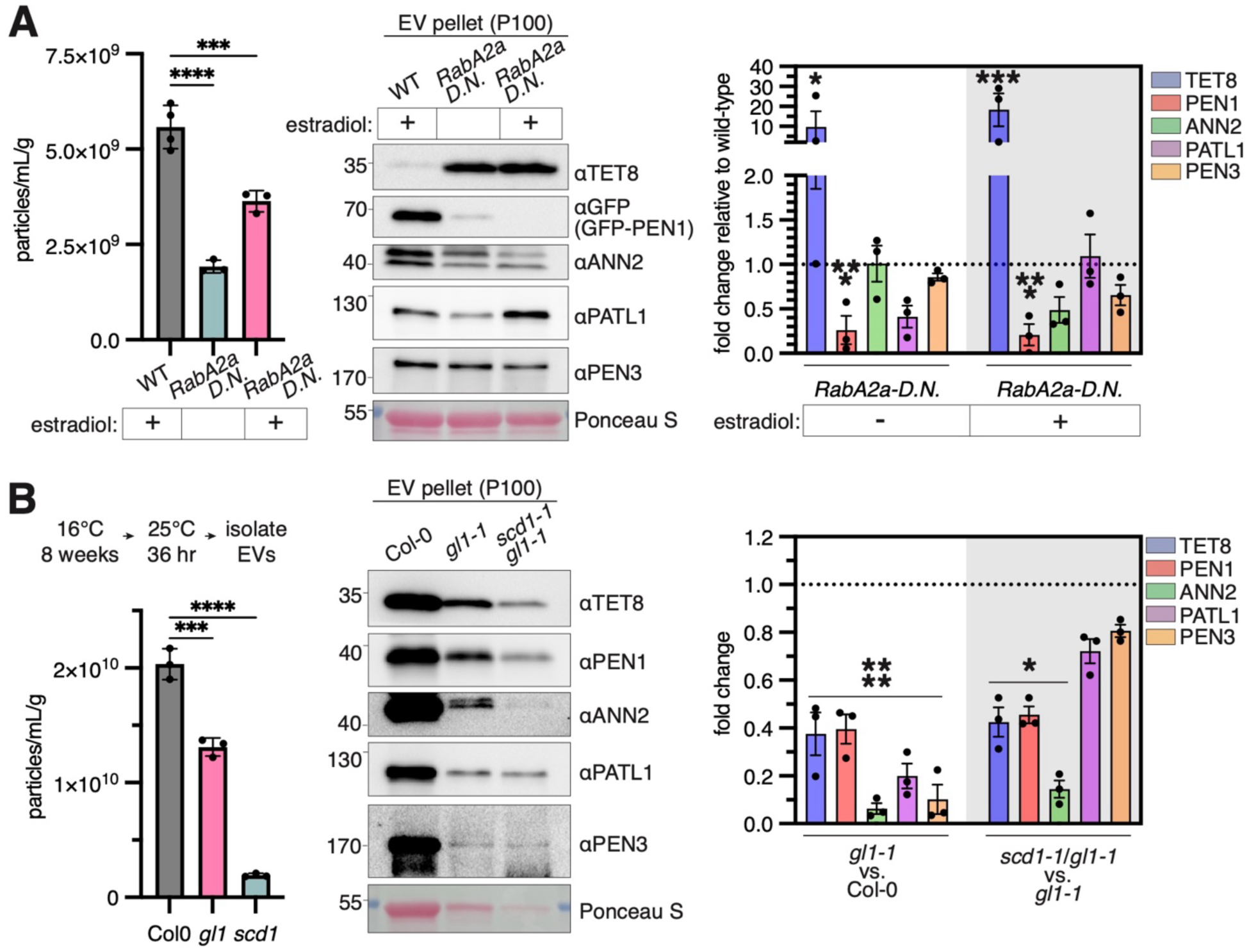
Rab proteins contribute to the secretion of EVs in a subpopulation-specific manner. NTA, immunoblot, and densitometry of EV pellets (P100) from wild-type and Arabidopsis mutants. Three independent biological replicates were collected for each genotype. For NTA, error bars indicate ± standard deviation, and statistical significance was detected using a one-way ANOVA and a Tukey post-hoc test, with p-value of <0.001 shown by ***; <0.0001 by ****; ns is not significant. For immunoblot, EVs collected from equivalent amounts of AWF were loaded in each lane. A representative immunoblot from 3 independent biological replicates is shown. Specificity of antibodies has been validated previously (Rutter and Innes, 2017; Koch et al. 2025). Densitometry was calculated in Fiji and error bars indicate ± SEM. Statistical significance for densitometry was detected by a two-way ANOVA with p-value of <0.05 shown by *; <0.0001 by ****. Biological replicate blots used for densitometry calculations are shown in Supplemental Fig. S7A. **A**) 10 µM estradiol or mock was applied by foliar spray 24 h prior to EV isolation. **B**) Plants were moved from 16°C to 25°C 36 h prior to EV isolation, a non-permissive temperature for the *scd1-1* allele, a GEF for RabE family GTPases. Fold change graph shows fold change of *gl1* mutant relative wild-type and fold change of *gl1scd1* double mutant relative to *gl1* single mutant.

We also utilized a temperature sensitive allele for SCD1, a Rab guanine nucleotide exchange factor (GEF) known to regulate the RabE family (Falbel et al., 2003; Mayers et al., 2017). When this mutant was shifted to a non-permissive temperature (26°C), the secretion of particles was significantly reduced (Fig. 7B). Immunoblotting showed that disruption of RabE family GTPases decreased the secretion of PEN1+, TET8+ and ANN2+ EV subpopulations (Fig. 7B). However, the Arabidopsis line carrying the *scd1-1* allele also carries a mutation in *GLABROUS1*, which encodes a Myb transcription factor required for formation of trichomes. We thus also analyzed a *gl1-1* single mutant. Unexpectedly, the *gl1-1* mutation by itself caused a large reduction in secretion of all EV markers tested (Fig. 7B). Because *GL1* encodes a transcription factor, it likely affects multiple cellular processes, thus it is unclear whether the reduction in EV secretion is a direct consequence of trichome loss, or if these are independent phenotypes caused by the loss of *GL1.* Notably, however, the *scd1gl1* double mutant showed a further reduction in TET8, PEN1, and ANN2 EV markers compared to the *gl1* single mutant (Fig. 7B), indicating that RabE family GTPases contribute to secretion of at least some EV subpopulations.

To test the hypothesis that RIN4 and EXO70 family proteins are involved in the secretion of multiple EV subpopulations, we isolated EV pellets (P100) from *rin4* and *exo70* mutant Arabidopsis plants. For RIN4, we used the *rin4/rps2/rpm1* mutant since mutation of *RIN4* by itself is lethal, but can be rescued by mutation of *RPS2* and *RPM1*, which are two disease resistance proteins that trigger cell death in the absence of RIN4 (Mackey et al., 2002). For EXO70 family proteins, we used *exo70a1, exo70b1, exo70b2, exo70e1, exo70e2, and exo70h1* (Chong et al., 2010; Li et al., 2010). With the exception of *exo70b1* and *exo70h1,* all of these mutant lines secreted fewer particles (Fig. 8). To test which EV subpopulations were responsible for this reduction in particle secretion, we probed the EV pellets of the *exo70* family and *rin4/rps2/rpm1* for a panel of EV marker proteins (Fig. 8, Supplemental Fig. S7B). In general, these mutants were depleted of PEN1+ and TET8+ EV subpopulations, with *rin4/rps2/rpm1, exo70e1,* and *exo70e2* mutants exhibiting the strongest depletion. ANN2+ EVs were specifically depleted from the *rin4/rps2/rpm1* mutant, while PATL1+ EVs were specifically depleted from most *exo70* family mutants. PEN3+ EVs were uniquely depleted in the *exo70a1* mutant. On the other hand, *exo70b1 and exo70b2* mutants did not exhibit EV subpopulation depletion, and *exo70b1* even exhibited significant increases in TET8+ and PEN1+ EVs (Supplemental Fig. S8), suggesting that there is functional specialization within the *exo70* family with regards to EV secretion. These results suggest that *rin4* and *exo70* proteins are involved in the biogenesis and secretion of specific subpopulations of EVs, especially the MVB-derived subpopulations marked by TET8 and PEN1.

**Supplementary Figure S7.**
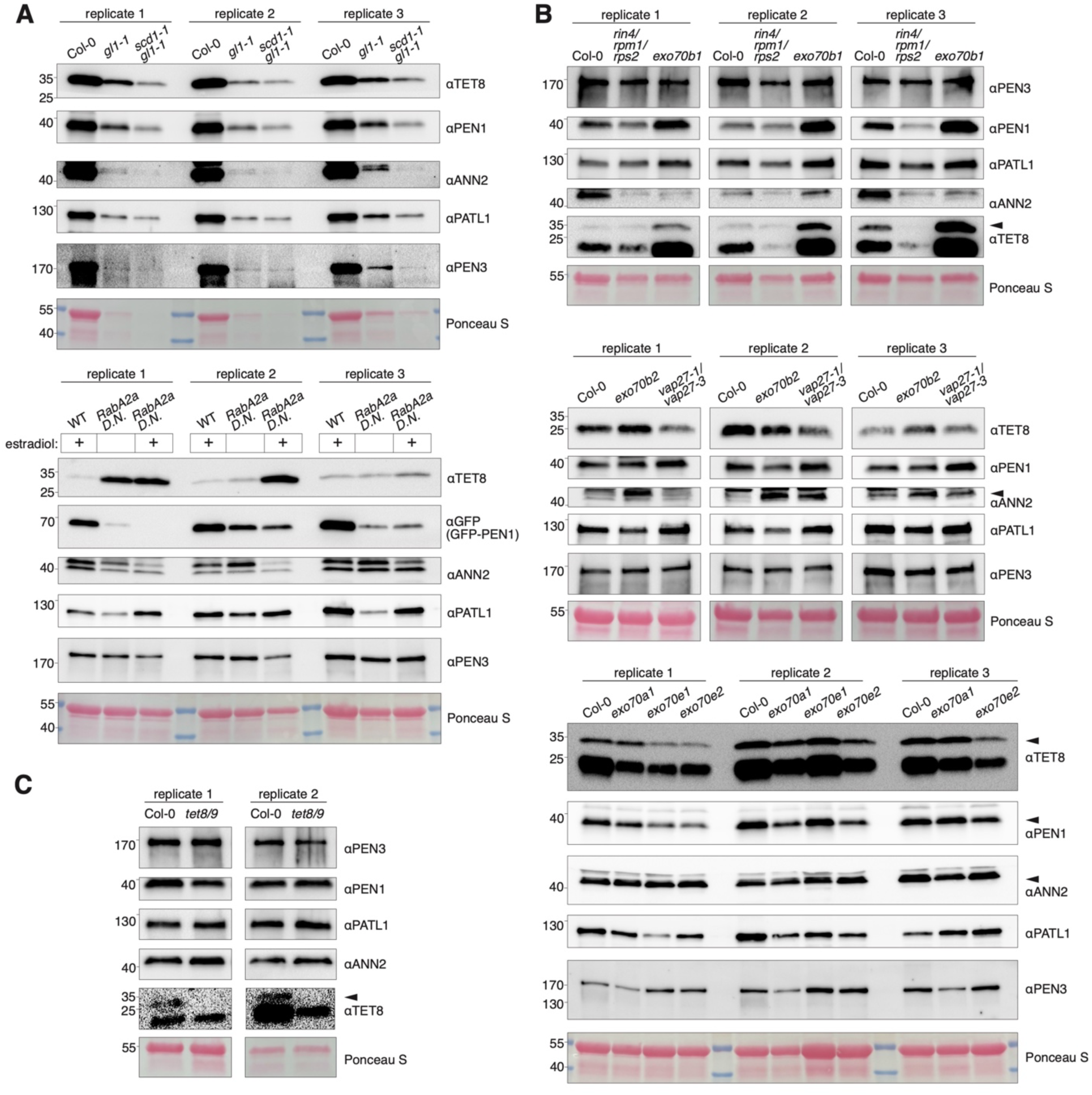
Identification of EV subpopulation secretion mutants by reverse mutant screen. Arrowheads indicate specific bands when other non-specific bands are present, such as for TET8 which has been previously described in (Koch, et. al. 2025). **A**) RabA2a and RabE family contribute to PEN1+ EV secretion and bulk EV secretion respectively. **B**) *exo70, rin4,* and *vap27* mutants lack specific EV subpopulations compared to wild-type (Col-0). *exo70e1* had only two biological replicates, so the average value was used for statistical analysis. **C**) TET8 does not contribute to the biogenesis of any other EV subpopulations that were tested. There were only two biological replicates, so the average value was used for statistical analysis.

**Figure 8.**
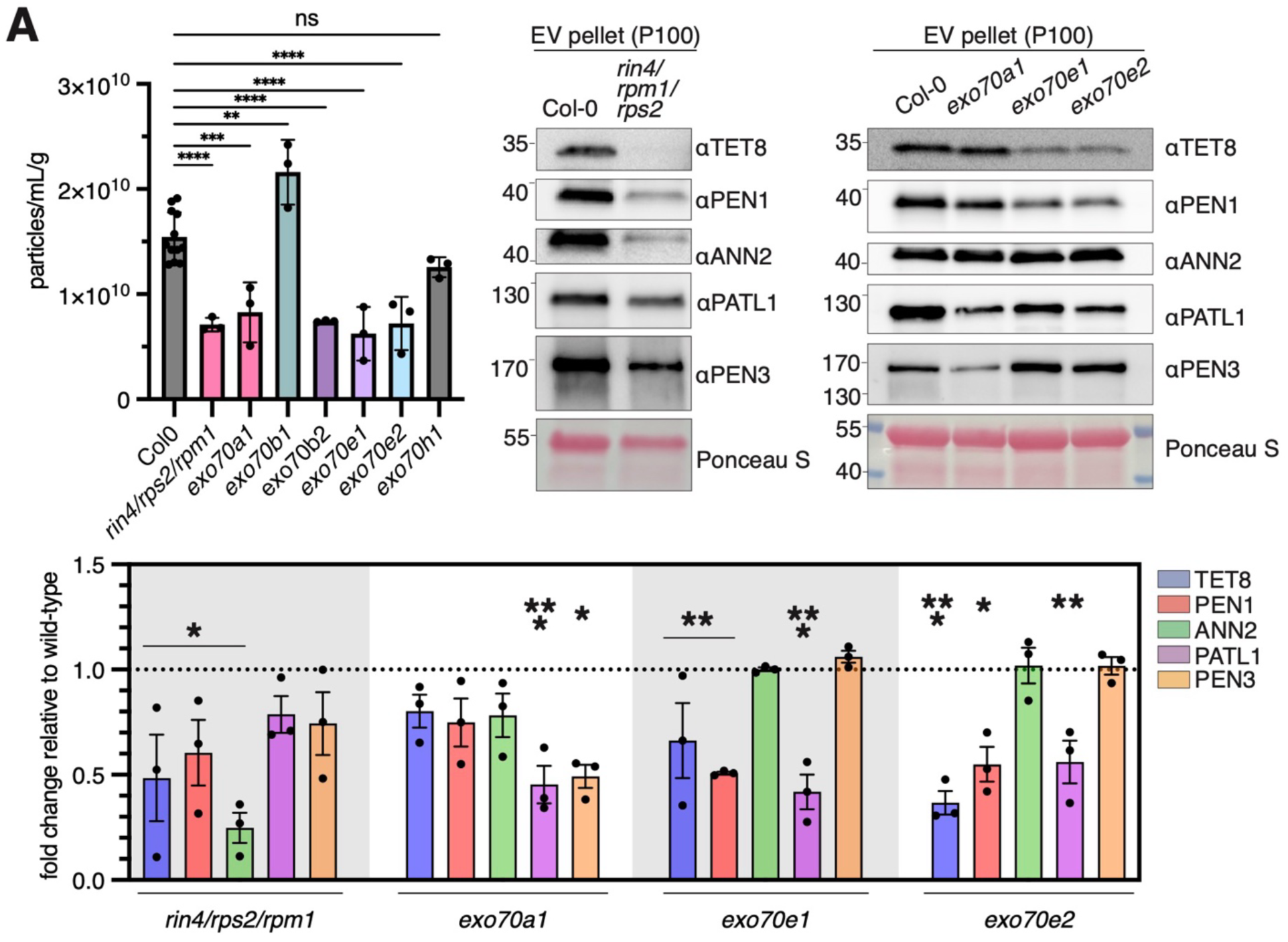
EXO70 family proteins contribute to the secretion of EV subpopulations. NTA, immunoblot, and densitometry of EV pellets (P100) from wild-type and Arabidopsis mutants. Three independent biological replicates were collected for each genotype. For NTA, error bars indicate ± standard deviation, and statistical significance was detected using a one-way ANOVA and a Tukey post-hoc test, with p-value of <0.01 is shown by **; <0.001 by ***; <0.0001 by ****; ns is not significant. *Exo70b1* had a premature senescence phenotype, likely accounting for its increase in particle secretion. For immunoblot, EVs collected from equivalent amounts of AWF were loaded in each lane. A representative immunoblot from 3 independent biological replicates is shown. For densitometry graph, error bars indicate ± SEM. Statistical significance was detected with a two-way ANOVA, followed by Dunnett’s post-hoc test, with p-value of <0.05 shown by *; <0.01 by **; <0.001 shown by ***; <0.0001 by ****. Biological replicate blots used for densitometry calculations are shown in Supplemental Fig. S7B.

**Supplementary Figure S8.**
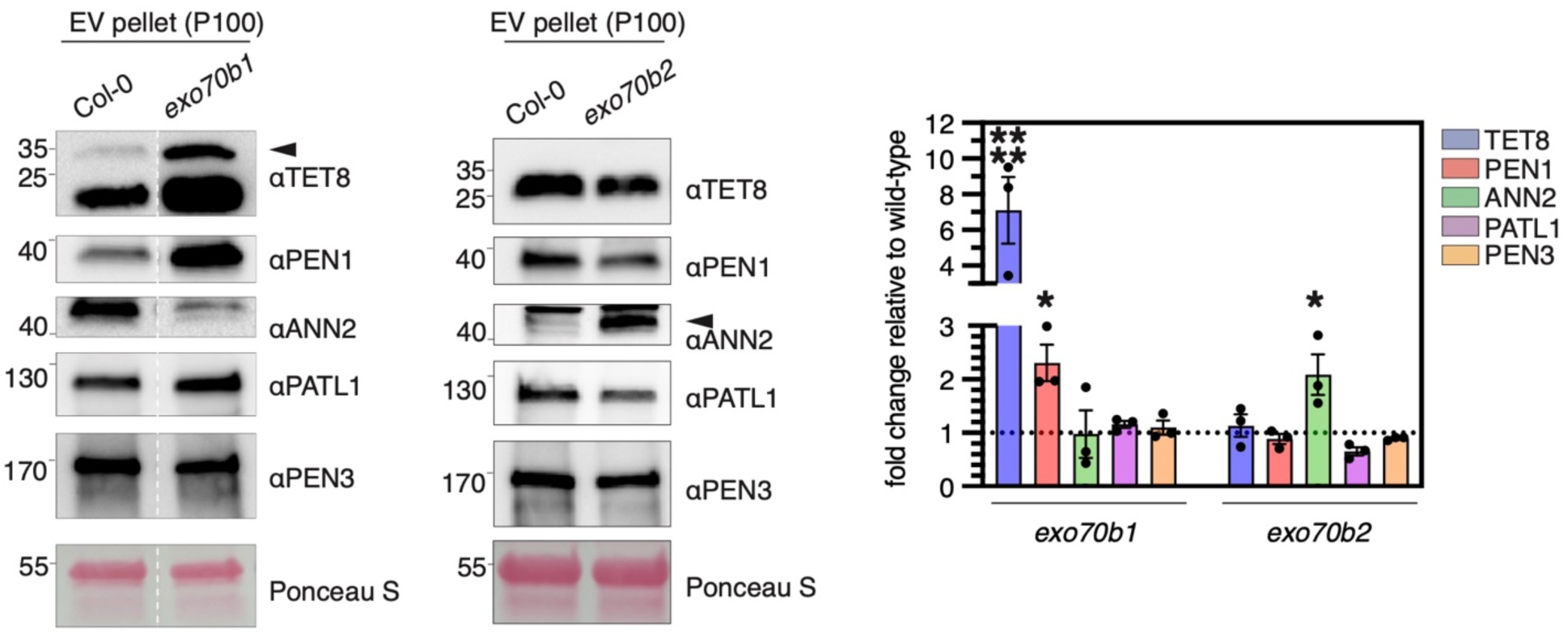
Neither *exo70b1* nor *exo70b2* are significantly impaired in EV secretion in mature plants. Immunoblot and densitometry of EV pellets (P100) from wild-type and Arabidopsis mutants. Three independent biological replicates were collected for each genotype. For densitometry graph, error bars indicate ± SEM. Statistical significance was detected with a two-way ANOVA, followed by Dunnett’s post-hoc test, with p-value of <0.05 shown by *; <0.0001 by ****. The *exo70b1* blot was stitched together from larger blots, and the dotted white line indicates where lanes were omitted from the original blots, which are shown in Supplemental Fig. S7A,C.

We noted that *rin4/rps2/rpm1* plants had a glabrous phenotype (Supplementary Fig. S9A). We therefore investigated the possibility that *gl1* was mutated in this line, but the sequence was wild type. We also noticed that *rin4/rps2/rpm1* had yellow seeds, suggesting that it may have a mutation in *ttg1,* a transcription factor involved in trichome development that also results in yellow seeds when mutant (Supplementary Fig. S9B). After sequencing, we confirmed that there was a point mutation resulting in a single amino acid substitution (L248P) in the *ttg1* allele of the *rin4/rps2/rpm1* line, which is likely the cause of the glabrous and yellow seed phenotype in *rin4/rps2/rpm1* (Supplementary Fig. S9C). Therefore, it is unclear whether the *rin4/rps2/rpm1* plants were deficient in EV secretion due to a *ttg1* mutation and glabrous phenotype, or due to a mutation in *rin4*, or both.

Previous studies have shown that *tet8tet9* double mutants have reduced particle secretion, and that TET8 is involved in biogenesis of EVs through interaction with COP1 at the Golgi apparatus (Cai et al., 2018; Liu et al., 2024). In agreement with these studies, we also observed a slight decrease in particle secretion in the *tet8tet9* double mutant (Fig. 9). However, we found that EV subpopulations marked by PEN1, ANN2, PATL1, or PEN3 were not affected in the *tet8/tet9* mutant (Fig. 9). This result suggests TET8 is involved in the biogenesis of a very specific class of EV. Given that we observed that TET8 interacted with VAP27 proteins in our proximity labeling study, we also assessed whether mutations in *VAP27* affected EV secretion. Specifically, we isolated EV pellets from a *vap27-1vap27-3* double mutant (Stefano et al., 2018). TET8 was the only EV subpopulation that was suppressed by VAP27 knock-down (Fig. 9). Together, these results indicate that TET8+ EVs have a unique biosynthetic pathway that involves VAP proteins.

**Supplementary Figure S9.**
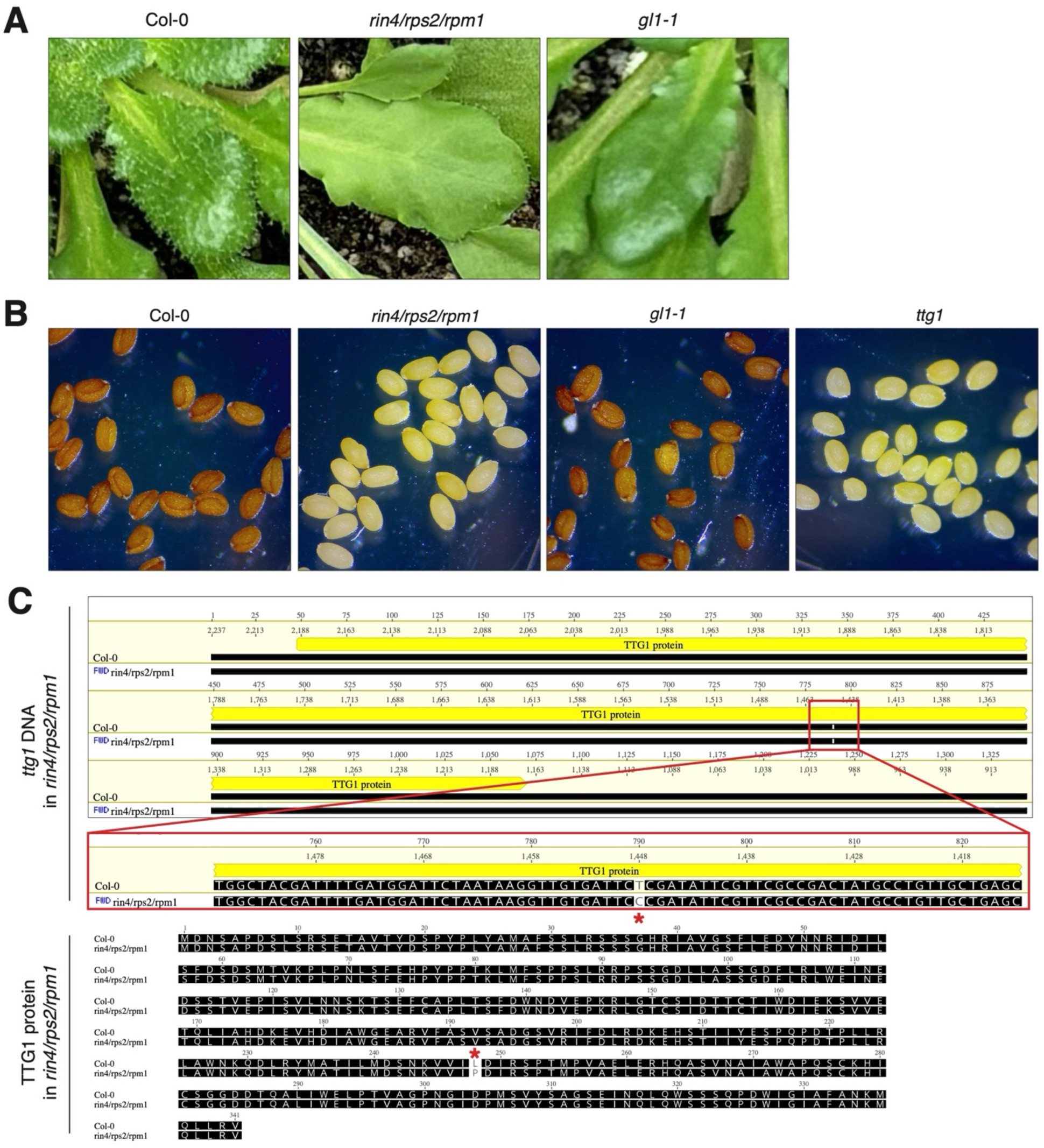
*rin4/rps2/rpm1* harbors a missense mutation in *ttg1,* a transcription factor involved in trichome development. **A)** Photographs of EV subpopulation mutant Arabidopsis lines, showing the glabrous phenotype of *rin4/rps2/rpm1,* compared to the glabrous *gl1-1* used in Fig. 7B. **B)** Micrographs of seeds showing yellow seed phenotype of *rin4/rps2/rpm1,* similar to that of *ttg1.* **C)** DNA and protein sequence alignments of *rin4/rps2/rpm1* compared to Col-0 at the *ttg1* locus. *rin4/rps2/rpm1* has a point mutation from T to C at position 790, resulting in a single amino acid substitution at position 248, indicated by red asterisks. This missense mutation likely explains the phenotypes in **A)** and **B)**.

**Figure 9.**
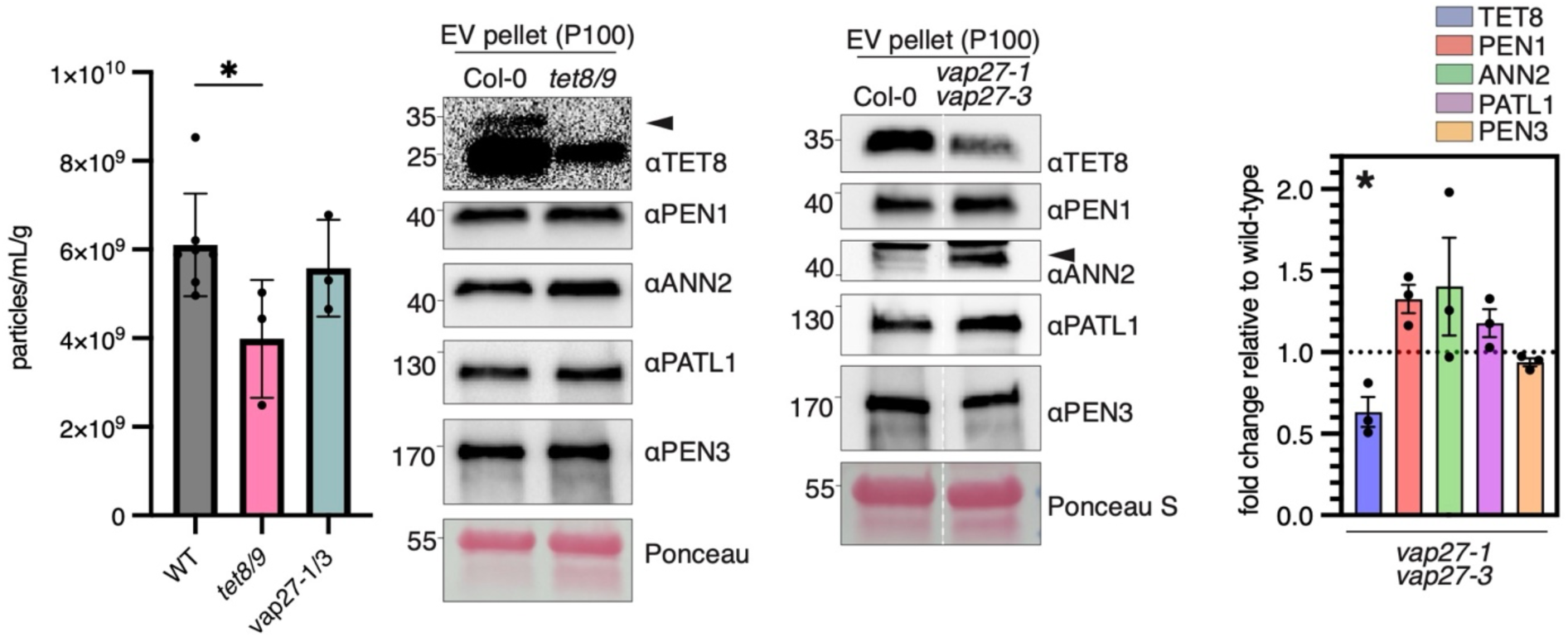
TET8 biogenesis is unique among EV markers tested. NTA, immunoblot, and densitometry of EV pellets (P100) from wild-type and Arabidopsis mutants. Three independent biological replicates were collected for each genotype. For NTA, error bars indicate ± standard deviation, and statistical significance was detected using a one-way ANOVA and a Tukey post-hoc test, with p-value of <0.05 shown by *. For immunoblot, EVs collected from equivalent amounts of AWF were loaded in each lane. A representative immunoblot from 3 independent biological replicates is shown. TET8-specific band is indicated with an arrowhead, and does not appear in the *vap27-1/vap27-3* blot because the membrane was cut to avoid probing the region with the non-specific band. For densitometry graph, error bars indicate ± SEM. Statistical significance was detected by two-way ANOVA and a Dunnett post-hoc test, with p-value of <0.05 shown by *. Additional lanes from the *vap27-1/vap27-3* blot were excised along the white dotted line. Original blots are shown in Supplemental Fig. S7C.

EVs have been proposed to contribute to plant immunity (Rutter and Innes, 2017; Cai et al., 2018; He et al., 2021; Koch et al., 2025). To test this prediction, we infected seedlings of EXO70 family mutants with the hemibiotrophic fungal pathogen *C. higginsianum* (*Ch*) and assessed infection progression. *Ch* infection begins with spore germination and appressoria formation on the leaf surface (0-48 hours post inoculation [hpi]). Penetration of the cell wall is followed by a short biotrophic phase in which short, bulky biotrophic hyphae feed on living plant cells (48-64 hpi). The necrotrophic phase is marked by extension of necrotrophic hyphae into adjacent cells and plant cell death (>64 hpi). We assessed *Ch* progression 54 hpi, just as *Ch* biotrophic hyphae were forming in wild-type seedlings. At this same time point, *Ch* infection of *exo70* family mutants had already progressed to late biotrophy/early necrotrophy (Fig. 10). All *exo70* family mutants had significantly more biotrophic hyphae. Both *exo70e1* and *exo70e2* mutants were particularly susceptible and had significantly more instance of necrotrophic hyphae than wild-type plants. Even *exo70b1* and *exo70b2* mutants were more susceptible, even though they did not have severe EV phenotypes (Fig. 10, Supplementary Fig. S8). This observation suggests that *EXO70b* proteins are involved in conventional post-Golgi secretion that contributes to fungal immunity independent of EVs. An alternative explanation is that while mature *exo70b* mutant plants did not have any EV defects, *exo70b* seedlings may indeed have EV secretion defects, and are thus more susceptible to *Ch* infection. Together, these results indicate that EXO70 family proteins contribute to the biogenesis and/or secretion of specific EV subpopulations that in turn contribute to resistance to fungal infection.

**Figure 10.**
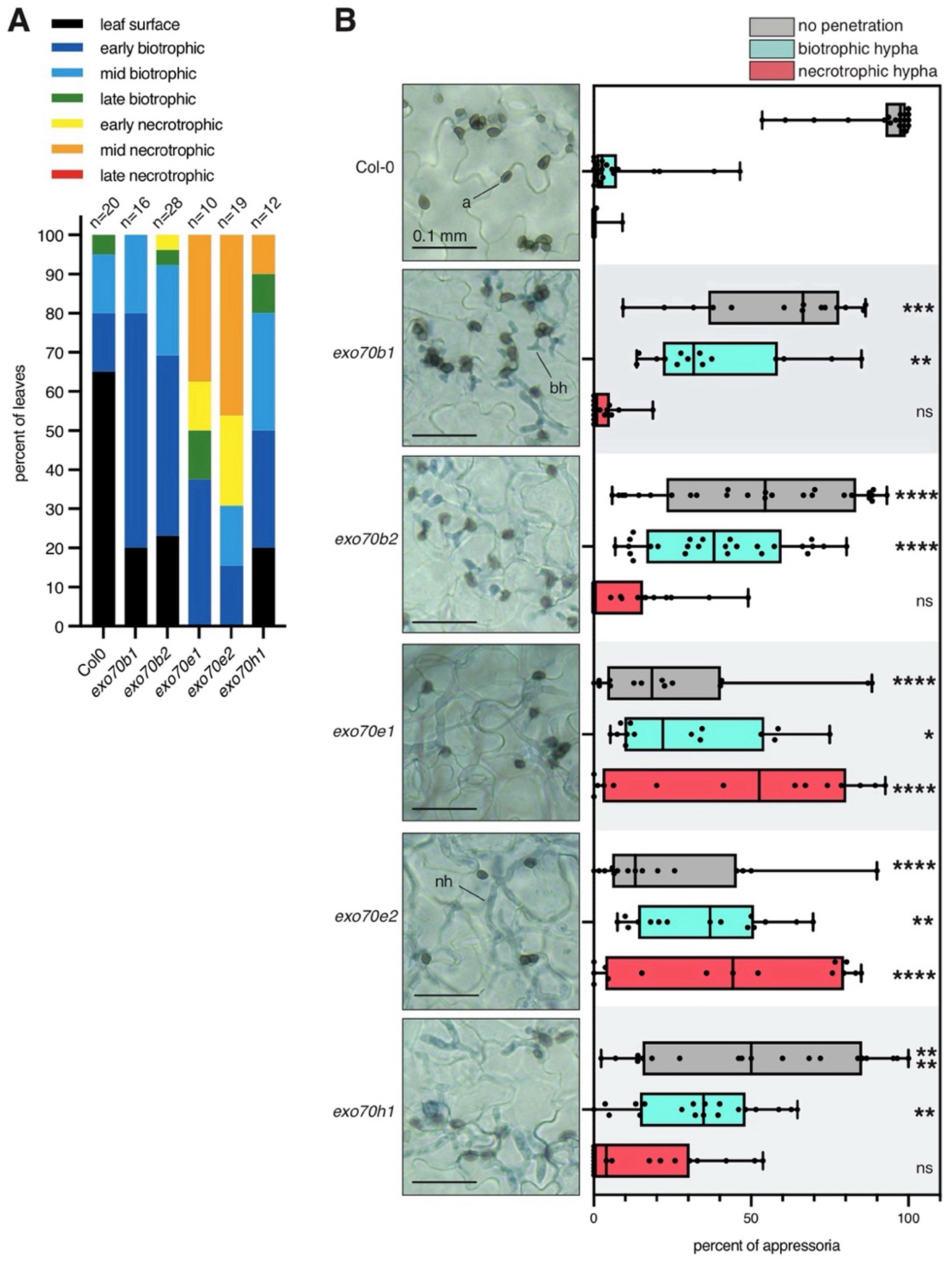
*exo70* family mutants are more susceptible to infection with *C. higginsianum.* Cotyledons of 10-day old seedlings were inoculated with 1000 spores of *C. higginsianum (Ch)* and assessed for fungal growth stage 54 hpi using trypan blue staining. **A)** Stage of infection was scored by assessment of biotrophic and necrotrophic hyphae. N indicates the number of cotyledons assessed per genotype. **B)** Images from a random location on each cotyledon were captured at 40X magnification, and events occurring at each appressoria (a) were manually categorized into no penetration, biotrophic hypha (bh), and necrotrophic hypha (nh). Each dot in the corresponding graphs represents the value from one cotyledon. Statistical significance was determined using a two-way ANOVA and post hoc Dunnett test, with with p-value of <0.05 shown by *; <0.01 shown by **; <0.001 by ***; <0.0001 by ****; ns not significant.

## Discussion

Our findings suggest a role for the exocyst complex in the secretion of EVs. The core exocyst complex is composed of eight subunits, one of which is EXO70 (exocyst component of 70 kDa) (Mei et al., 2018). In most other eukaryotes, including humans, EXO70 is encoded by a single gene, but in plants the EXO gene family is highly expanded, with the Arabidopsis genome containing 23 paralogs (Zarsky et al., 2009). This expansion suggests there may be multiple forms of the exocyst complex, each with a specialized function (Chong et al., 2010; Li et al., 2010; Markovic et al., 2021). Several EXO70 family proteins are implicated in pathogen resistance, including EXO70B1, EXO70B2, EXO70E1, and EXO70H1 (Pečenková et al., 2011; Ortmannová et al., 2022). Recently, we identified exocyst proteins, including EXO70A1, as proteins secreted with EVs (Koch et al., 2025). Here we describe a specific role for different EXO70 family proteins in secretion of specific EV subpopulations. EXO70E1 and EXO70E2 were the most essential of the EXO70 family proteins in secretion of immunity-related EV subpopulations marked by PEN1 and TET8, which possibly explains why *exo70e1 and exo70e2* mutants were the most susceptible to infection by *C. higginsianum* among the *exo70* family mutants we tested. Notably, EXO70E2 is essential for formation of the unique plant organelle EXPO, a double membraned organelle that fuses with the plasma membrane using exocyst components for the secretion of a single, large, intraluminal vesicle (Wang et al., 2010; Ding et al., 2014; Lin et al., 2015). Perhaps EXPO plays a role in EV secretion for pathogen defense, but further work is needed to test this hypothesis.

EXO70B1 contributes to papillae formation for powdery mildew defense, and EXO70B2 was found to localize to powdery mildew haustoria (Pečenková et al., 2011; Ortmannová et al., 2022). In the Ortmannová study, PEN1 was even shown to interact with EXO70B2. Despite these promising connections, we did not find a large effect on EV secretion in *exo70b1* and *exo70b2* mutants. Perhaps these *EXO70* paralogs have redundant function or are involved in secretion of other non-vesicular cargo for pathogen defense at sites of infection. Our study also found that *exo70a1* mutants were deficient in secretion of PEN3+ and PATL1+ EVs. PEN3 is an ABC transporter that localizes to sites of pathogen infection to transport antimicrobial metabolites such as camalexin, glucosinolates, and other compounds (Stein et al., 2006; He et al., 2019; Matern et al., 2019). EXO70 appears to play a role in EV secretion in other eukaryotes. In human cells, RNAi knockdown of EXO70 resulted in decreased secretion of several EV subpopulations (Liu et al., 2023). The exocyst complex was also found to localize with some but not all MVBs using TIRF microscopy (Liu et al., 2023). Our results indicate that the role of the exocyst complex in EV secretion may be conserved in plants.

Rab GTPases are another class of proteins whose role in EV secretion is widely conserved, even between Archaea and eukaryotes (Mills et al., 2024). Rabs are a large protein family, with the current count standing at about 60-70 Rabs in both humans and Arabidopsis (Homma et al., 2020; Tripathy et al., 2021). The secretion of RabA family proteins in EVs has been shown to be induced by infection with the fungal pathogen *C. higginsianum,* suggesting that RabA proteins might contribute to the secretion of specific EV subpopulations in response to biotic and abiotic stress (Koch et al., 2025). In this study, we present evidence that RabA2a is specifically involved in secretion of PEN1+ EVs, and SCD1 (a GEF for RabE’s) is involved in secretion of PEN1, TET8, and ANN2 EV subpopulations. A recent study found that RabA2a recruits PEN1 to the plasma membrane (Pang et al., 2022). Given that Arabidopsis EVs are enriched in plasma membrane proteins (Rutter and Innes, 2017), we speculate that PEN1+ EVs are derived from recycling of PM nanodomains. The reduction of PEN1+ EVs observed in the RabA2a dominant negative line may thus be due to the general reduction in plasma membrane localized PEN1. Our results also indicate that RabA2a and PEN1 function is not essential for the secretion of other EV subpopulations. This is surprising given that PEN1 is co-secreted with PEN3 in >40% of EVs (Koch et al., 2025). However, the Pang study also showed that PEN3 can be recruited to the plasma membrane by exocyst components independent of PEN1 and RabA2a (Pang et al., 2022). This suggests that parallel pathways exist for the secretion of different EV subpopulations. Still, PEN1 and TET8 seem to localize to the same MVB-like subcellular structure, with the caveat that this could be due to the overexpression of these proteins in our system. Indeed, our results indicate that RabE family members are important for the secretion of several EV subpopulations, including both PEN1+ and TET8+ EVs. Notably, the human ortholog of Arabidopsis RabE1b is specifically involved in secretion of CD63, a human homolog of TET8 (Speth et al., 2009; Temoche-Diaz et al., 2019).

RIN4 is vital to plant immunity and stands at the center of the plant-pathogen arms race, where the two guard proteins RPM1 and RPS2 monitor its targeting by pathogen effectors (Mackey et al., 2002; van der Hoorn and Kamoun, 2008). Despite its importance in immunity, the function of RIN4 itself was unknown. Here we have presented data that support a general role for RIN4 in the biogenesis and secretion of EVs, which would represent a major step in our understanding of RIN4. We found that RIN4 interacted with EV marker proteins, and that RIN4 localized to the outer surface of MVB-like subcellular structures. A recent study showed that RIN4 has a prominent intrinsically disordered region (IDR) that enables liquid-liquid phase separation (LLPS) of RIN4 in vitro and in vivo (Zhu et al., 2025), which could conceivably contribute to packaging of RIN4 into EVs. Two other recent reports in plants have suggested that LLPS may have a role in vesicle trafficking. One study showed that a plant ESCRT-associated protein FREE1 forms condensates in vitro and in vivo on the surface of MVBs (Wang et al., 2024). When condensate formation of FREE1 was induced in vivo, a significant enough force was generated to produce intraluminal vesicles (ILVs), suggesting that LLPS alone may be sufficient for formation of ILVs on MVBs (Wang et al., 2024). Another study showed that the ESCRT-associated protein ALIX and a lipid binding protein form condensates in response to Ca^2+^ (Mosesso et al., 2024). FREE1 and ALIX both contain IDRs that are required for LLPS and condensate formation, therefore it would be interesting to know whether RIN4 also falls into this emerging category of proteins that localize to MVBs via formation of biomolecular condensates. One important caveat of our current experiments is that it remains unclear whether *rin4/rps2/rpm1* is deficient in secretion of EVs due to the *rin4* mutation or to a mutation in the transcription factor *TGG1*, which rendered these plants glabrous. Future work will need to assess whether the *rin4/rps2/rpm1* triple mutant displays reduced EV secretion in a wild-type *TTG1* background.

Trichomes vary among plants but are generally involved in protection from abiotic and biotic stress. Arabidopsis has relatively simple, unicellular, non-glandular trichomes (Schuurink and Tissier, 2019). It is tempting to hypothesize that trichomes might be a hot spot for EV secretion. Indeed, a recent study of Arabidopsis trichomes found that numerous EV marker proteins are among the highest expressed proteins in trichomes, including PEN1, TET8, ANN2, PEN3, PATL1, and many Rab protein and V-ATPases common in the EV proteome (Huebbers et al., 2022; Huebbers et al., 2024; Koch et al., 2025). Trichomes also function in mechanosensation, and perhaps mechanosensation generated by trichomes is required to stimulate EV release (Matsumura et al., 2022). In this study, we found that mutation of *GL1* was sufficient to reduce the secretion of most EV subpopulation markers by 50%. Though a simple explanation for this effect is that trichomes are hot spots for EV secretion, GL1 is a MYB-like transcription factor that works together with several other transcription factors including TTG1 in a complex network to help differentiate epidermal cells for trichome development (Balkunde et al., 2020). As such, GL1 likely affects the expression of numerous target genes, and the effect of *gl1* mutation on those downstream targets may have led to a decrease in EV secretion.

Perhaps more compelling is the observation that PEN1 is highly expressed in guard cells and PEN1 MVB-like structures are readily observed in guard cells, suggesting that guard cells may be a source of PEN1+ EV secretion. Guard cells are important for plant immunity, because guard cells regulate stomatal aperture during infection, acting to restrict the access of pathogenic bacteria and fungi to the apoplast (Melotto et al., 2006; Wu and Liu, 2022). We speculate that PEN1+ EVs are specifically secreted by guard cells as this is where plant EVs will most likely encounter a plant pathogen. Future studies of plant EVs should more thoroughly investigate which cell types contribute to the secretion of distinct EV subpopulations.

One possibility we assessed in this study is that autophagosomes could be one source of EVs. Autophagy proteins and autophagosomes have been observed to localize to sites of pathogen attack, and autophagic flux is increased during infection (An et al., 2006; Dagdas et al., 2018; Pandey et al., 2021; Regmi et al., 2024). In animal cells, autophagy coordinates with the endomembrane system for secretion of autophagy substrates including ATG8 (also called LC-III in humans) through the formation of hybrid organelles called amphisomes (Guo et al., 2017; Jeppesen et al., 2019; Leidal et al., 2020). However, we have been unable to identify ATG8 proteins in EVs purified from healthy or *C. higginsianum-*infected Arabidopsis (Koch et al., 2025), and we failed to observe co-localization of ATG8a with EV marker proteins. Despite these data, we found that mutations in *ATG2* and *ATG5* strongly suppressed secretion of TET8+ EVs, suggesting that autophagy may indirectly contribute to secretion of a specific subset of EVs.

In conclusion, in this study we used an unbiased proximity labeling approach, which helped identify several proteins and pathways involved in the secretion of plant EV subpopulations, including EXO70, RIN4, and RabA and RabE family proteins. Understanding the biogenesis of EV subpopulations is important given that plant EV subpopulations may play distinct roles during pathogen infection. We hope that this study lays the groundwork for future investigation of the biogenesis of plant EVs.

## Materials and methods

### Plant materials

The Arabidopsis genotypes are indicated in Supplementary Table S1. Arabidopsis plants were grown in Sungro Propagation mix in a growth room at 22°C-24°C, 40%-60% relative humidity, and under 150 *µ*mol m^-2^ s^-1^ light for 9 h per day. Illumination was provided by GE HI-LUMEN XL Starcoat 32-watt fluorescent bulbs (a 50:50 mixture of 3,500 and 5,000 K spectrum bulbs). Miracle-Gro water-soluble fertilizer (catalog no. 24-8-16) was used to water the plants every 10-15 days (250 mg/L).

**Supplemenatary Table S1.**
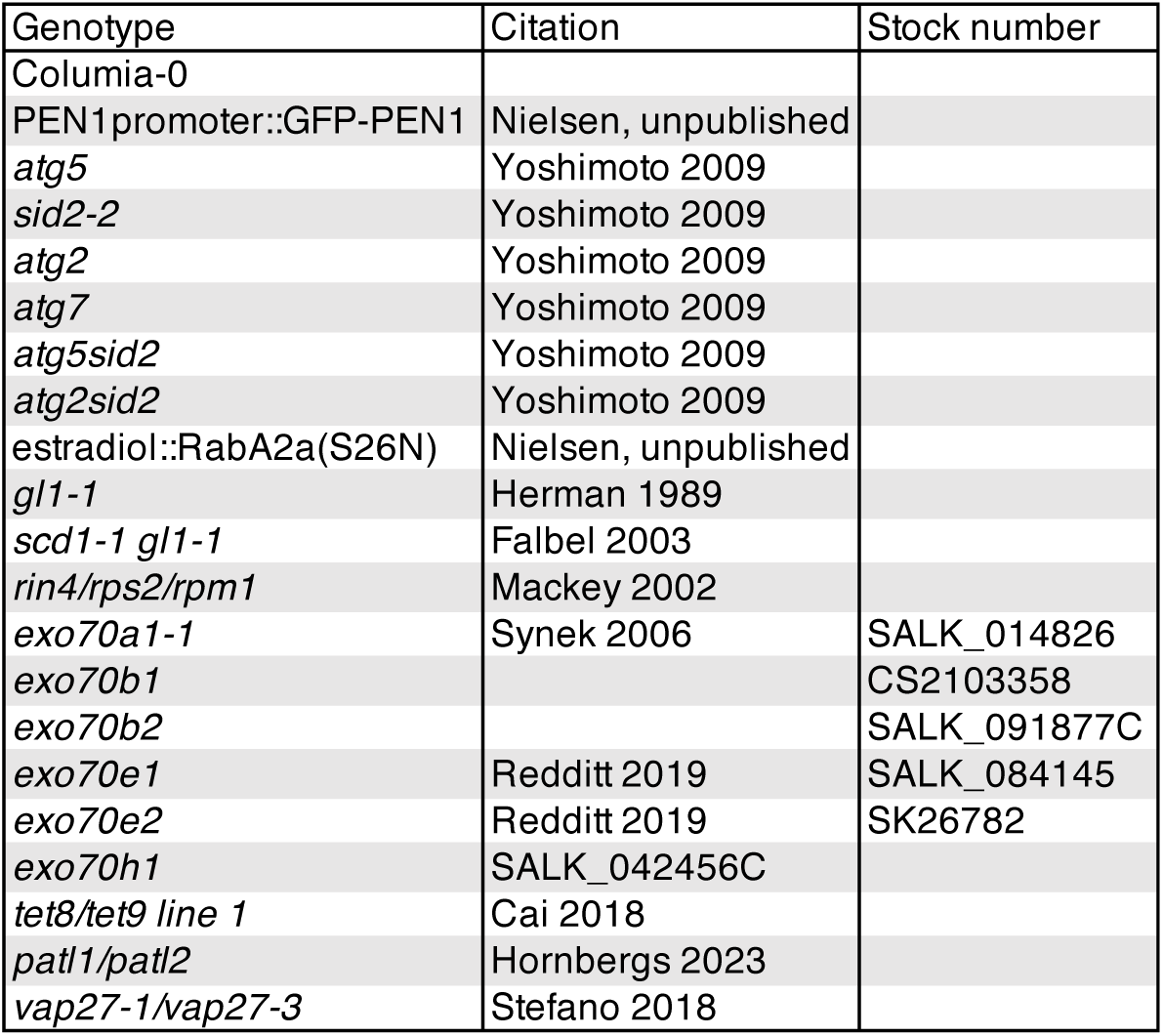
Arabidopsis lines used in this study.

### Isolation of apoplastic wash fluid

Apoplastic wash fluid (AWF) was isolated from Arabidopsis using a previously published protocol described in Koch et al. (2025) adapted from that described by Rutter et al. (2017). To isolate EVs from *N. benthamiana,* leaves of 3-4-week-old *N. benthamiana* were detached, washed in distilled water three times, and infiltrated by placing in a beaker abaxial side facing up, submerging in 4°C vesicle isolation buffer supplemented with EGTA (VIB, 10mM EGTA, 20 mM MES, 2 mM CaCl_2_, and 10 mM NaCl, pH 6) using a French press coffee maker. EGTA was included to reduce co-pelleting of polysaccharides from *N. benthamiana* AWF, which formed a gel that interfered with pelleting of *N. benthamiana* EVs. Submerged leaves and rosettes were placed in a vacuum chamber and a vacuum applied for 20 sec using a vacuum pump (Thermo Savant VP 100 Two Stage Model #1102180403) until ∼95% of leaf area appeared infiltrated. Infiltrated leaves and rosettes were patted dry with paper towel to remove all excess buffer. Leaves and rosettes were rolled and placed stem cut site facing up inside modified 50 mL syringes with 10-20 extra holes melted into the bottom to allow buffer to freely flow through. The loaded syringes were placed in 250 mL wide-mouth bottles using parafilm wrapped around the syringe as an adaptor where necessary to hold the syringe in the bottle mouth. Bottles were placed in a fixed-angle rotor (Beckman JA-14 #339247) and centrifuged at 500 × *g* for 20 min at 4°C using slow acceleration to isolate AWF from the leaves. AWF was passed through a 0.45 *µ*m filter (PALL Life Sciences #4612, EMD Millipore #SCGP00525).

### EV pellet isolation

EV pellets were isolated from AWF as previously described in Koch et al. 2025. For differential ultracentrifugation of *N. benthamiana* AWF in Fig. 1, filtered AWF was centrifuged at 10,000 × *g* for 30 min at 4°C (P10). The supernatant was transferred to a new tube and centrifuged at 40,000 × *g* for 1 h at 4°C (P40). The supernatant was decanted to a new tube and centrifuged at 100,000 × *g* for 1 h at 4°C (P100). Pellets were resuspended in filter-sterilized 20 mM Tris HCl pH=7.5. For Arabidopsis experiments, where a P100 was used, the P10 supernatant was centrifuged at 100,000 × *g* for 1 h at 4°C and resuspended as before – no 40,000 × *g* spin was performed.

### Enzymatic digestion of EV pellets

For protease protection assays, EV pellets were resuspended in ultraclean 150 mM Tris HCl pH=7.8 buffer and aliquoted. EV pellets were used immediately when performing protease protection assays. For detergent treatment, 6% v/v Triton X-100 (EMD Millipore #TX1568-1) detergent stock was added for a final concentration of 1% TX-100, and the sample was incubated for 30 min at 4°C. For protease treatment, a 10 *µ*g/mL stock of trypsin (Promega #V5113) was added to a final concentration of 1 *µ*g/mL, and the sample was incubated for 1 h 37°C.

### Nanoparticle tracking analysis (NTA) and transmission electron microscopy (TEM)

Resuspended EV pellets were used immediately (without freezing) for NTA or TEM analysis. For NTA, the resuspended pellets were diluted in ultraclean Tris resuspension buffer. The nanoparticle tracker (Particle Metrix PMX 120 Zetaview Mono 488 Laser running Zetaview software version 8.05.16 SP3) was calibrated with 100 nm beads (nanoComposix silica nanospheres #SISN100). Diluted EV samples (ranging from 1:500 to 1:2000 dilutions) were injected such that approximately 100 particles were visible per field of view and could be observed when using the camera settings of 75 sensitivity, 30 frame rate, and 100 shutter. Particle concentration was recorded as the average of 11 camera positions. For TEM, copper or nickel formvar-coated grids (Electron Microscopy Sciences, # FCF300) were prepared using glow discharge at 0.4 mBar and 15 mA for 1 min (PELCO easiGlow #91000 and #91040). 5 *µ*L of each sample was pipetted onto the formvar-coated surface and allowed to adhere for 5 min. Sample was wicked from the grid using Whatman filter paper and stained twice with 10 *µ*L 2% uranyl acetate by pipetting the uranyl acetate across the surface of the grid. Stained grids were protected from light and allowed to dry at least 16 h before imaging at 80kV using a JEOL JEM-1010 transmission electron microscope.

### Protein preparation and immunoblots

Leaves were flash-frozen in liquid nitrogen and ground into a fine powder. Cold protein extraction buffer (RIPA, 500 mM NaCl, 50 mM Tris HCl pH 7.5, 1mM EDTA, 1% v/v Nonidet-P40, 0.5% w/v sodium deoxycholate, 0.1% SDS, 1% v/v plant protease inhibitor cocktail (Millipore Sigma catalog number P9599), 1mM DTT) was added to equal masses of ground tissue and mixed with the sample on ice using a mechanical pestle for 1 min. After incubation on ice for 20 min, debris was pelleted twice by centrifuging at 16,000 × *g* for 10 min at 4°C.

SDS-PAGE and immunoblotting were performed as previously described (Koch et al. 2025). Briefly, samples from either equal masses of lysate or EV pellets collected from equivalent volumes of AWF were mixed with 5X loading dye (250 mM Tris HCl pH 6.8, 10% w/v SDS, 40% v/v glycerol, 0.02% bromoethanol blue, 5% v/v 2-mercaptoethanol), incubated at 95°C for 5 min, and run in 4–20% TGX stain-free denaturing protein gels (Biorad #4568093, #4568096, #5678094, #5678095). Protein was transferred to 0.45 *µ*m nitrocellulose membrane (Cytiva Amersham Protran #10600003) and visualized by applying Ponceau Stain (0.1% w/v Ponceau S Stain, 5% v/v acetic acid) for 3 min and rinsing in distilled water.

For immunoblot, membranes were blocked in 5% skim milk dissolved in Tris-buffered saline + Tween 20 (TBST, 100 mM Tris, 150 mM NaCl, 0.1% Tween-20, pH 7.5) for 1 h at 25°C with gentle rocking – with the exception of streptavidin-HRP blots, which were blocked in 2.5% BSA, since milk inhibits the binding of streptavidin to biotin (Hoffman 1989). After blocking, membranes were incubated with primary antibody for 16 h at 4°C, washed (see Supplementary Table S2), incubated with secondary antibody where applicable, washed, and incubated with enhanced chemiluminescent substrate (National Diagnostics Protoglow #CL-300) before detecting signal using a Chemidoc MP imaging system (Biorad #12003154) or by exposing membrane to film, developed using a table-top x-ray film processer (Konica #SRX101A). Densitometry was calculated using FIJI. Graphs and statistical analyses were produced in Graphpad Prism. The specificity of antibodies was confirmed by probing Arabidopsis null mutants (Koch et al. 2025).

**Supplementary Table S2.** Antibodies used in this study.

| Antibody | Species | Dilution | Source | Catalog number |
| --- | --- | --- | --- | --- |
| PENETRATION 1 (PEN1) | Rabbit | 1:1000 | Zhang et. al. 2007 | N/A |
| PATELLIN 1 (PATL1) | Rabbit | 1:2000 | Peterman et. al. 2004 | N/A |
| TETRASPANIN 8 (TET8) | Rabbit | 1:800 | PhytoAB | PHY1490A |
| RPM-1 INTERACTING PROTEIN 4 (RIN4) | Rabbit | 1:2000 | Belkhadir et. al. 2004 | N/A |
| PENETRATION 3 (PEN3) | Rabbit | 1:1000 | Agrisera | AS224846 |
| ANNEXIN 2 (ANN2) | Rabbit | 1:1000 | PhytoAB | PHY1021A |
| GFP | Mouse | 1:2000 | Sigma-Aldrich | 66002-1-Ig |
| V5 | Mouse | 1:1000 | ChromoTek | (SV5-P-K) |
| myc-HRP | Mouse | 1:1000 | Abcam | [9E10] |
| streptavidin-HRP (use BSA not milk) |  | 1:10,000 | Abcam | ab7403 |
| Rabbit-HRP | Goat | 1:10,000 | Abcam | AB97051 |
| Mouse-HRP | Goat | 1:10,000 | Abcam | AB6789 |

### Affinity purification

Biotinylated proteins were affinity-purified using streptavidin-conjugated magnetic beads as previously described (Dinesh-Kumar et al., 2020). Briefly, protein samples were desalted using Zeba™ Spin columns (Thermo #89893) to remove excess biotin. Protein concentration was determined using a Bradford assay and BSA standard curve, and equal amounts of protein (2-4 mg depending on the experiment) were mixed with 100 *µ*L streptavidin beads (Dynabeads™ MyOne™ Streptavidin C1, Thermo #65001). After incubation, beads were washed sequentially 10 times for 8 min each in various buffers (Dinesh-Kumar et al., 2020) and collected by magnetic rack (Promega) between washes. A small aliquot of beads was mixed with 5X SDS loading dye including excess biotin, and the rest of the dry beads were flash frozen until analysis by semi-quantitative mass spectrometry.

### Confocal fluorescence microscopy

Confocal fluorescence microscopy analyses were performed using both Leica Stellaris 8 and Leica SP8 microscopes fitted with 63x oil or water immersion objectives. To image GFP, 488 nm light was used to excite, and signal was detected from 498-550 nm; for mScarlet, 569 nm light was used to excite, and signal was detected from 579-620 nm; for mCherry, 587 nm light was used to excite, and signal was detected from 597-620 nm; for YFP, 515nm light was used to excite, and signal was detected from 525-577 nm (YFP was only used in combination with mCherry). All images except autophagosome images are from single planes. Images were processed in Fiji, with negative controls processed in the same way as positive controls. For Pearson correlation coefficient analysis, cells expressing both markers equally were selected as a region of interest (ROI) and analyzed using the JaCoP plugin. For line intensity plots, intensity values were found in Fiji and graphed in GraphPad Prism.

### Co-immunoprecipitation

Protein was extracted as described above from equal amounts of tissue (approximately 1g) but using a milder extraction buffer (coIP extraction buffer, 150 mM NaCl, 25 mM Tris HCl pH 7.5, 1 mM EDTA, 10% v/v glycerol, 1% v/v plant protease inhibitor cocktail, 10 mM DTT, 1 mM PMSF). Equal amounts of protein were added to 20 *µ*L GFP-TRAP agarose beads (Chromotek #gt-20). After 3 h incubation at 4°C, beads were washed 5 times in coIP extraction buffer with 0.1% v/v Nonidet-P40 (tube was inverted 5 times and beads were collected by centrifugation at 2500 × *g* for 5 min for each wash), resuspended in 5X SDS loading buffer, and boiled at 95°C for 5 min prior SDS-PAGE and immunoblot analysis.

### Plasmid construction and sequencing

Constructs were made using Multisite Gateway cloning (Thermo) and verified by whole-plasmid sequencing (Eurofins), as previously described (Margets et al., 2024). TET8 was amplified from cDNA (Verso cDNA Synthesis Kit, Thermo #AB1453A) with attB1 and attB4 sites and integrated into pBSDONR P1-P4 (Qi et al., 2012) using BP clonase (Thermo). ANN2 and VAP27-1 were amplified from Arabidopsis Gateway cDNA clones obtained from The Arabidopsis Biological Resource Center and integrated into pBSDONR P1-P4 as above. mScarlet was amplified from pUC57 (a gift from Tim Cioffi) with attB4 and attB2 sites and integrated into pBSDONR P1-P4. Myoxin XI-K globular tail domain (GTD) (Zhang et al., 2021) was amplified from pGEX (a gift from Chris Staiger) with attB4 and attB2 sites. For proximity labeling constructs, TET8-miniTurbo, ANN2-miniTurbo, and miniTurbo-PEN1 were assembled into pBAV154 (dexamethasone-inducible; (Vinatzer et al., 2006)) using LR clonase (Margets et al., 2024). For microscopy and coIP constructs, GFP-PEN1, ANN2-GFP, TET8-GFP, TET8-mCherry, mScarlet-RIN4, mCherry-EXO70B2, myc-PEN1, ANN2-myc, TET8-myc, and myc-EX70B2 were assembled into pTA7001-DEST (DEX-inducible; Vinatzer et al. 2006) using LR clonase. PEN1 pBSDONR P4r-P2, YFP-miniTurbo pBAV154, RIN4 pBSDONR P4r-P2, mCherry pBSDONR P4r-P2, 5xMyc pBSDONR P1-P4, 5xMyc pBSDONR P4r-P2, AvrPphB-GFP pTA7001, ARA6-mCherry pEG100, YFP-ATG8a pEG100, and miniTurbo multisite Gateway clones were made as previously described (Gu and Innes, 2012; Rutter and Innes, 2017; Redditt et al., 2019; Margets et al., 2024). For estradiol inducible RabA2a-NI, the cDNA encoding RabA2a-N125I was amplified from a previously established line (Chow et al., 2008) and subcloned into pENTR-D-TOPO. RabA2a-NI was introduced into a modified pMDC7 vector (Lee et al., 2013). All primers used for cloning are listed in Supplementary 3S3.

The *TTG1* and *GL1* loci were sequenced by amplifying these genes using Phusion polymerase (ThermoFisher) using Arabidopsis genomic DNA isolated from Col-0 and *rin4/rps2/rpm1* mature leaf tissue as template. The primer sequences are listed in Supplementary Table S3.

**Supplementary Table S3.**
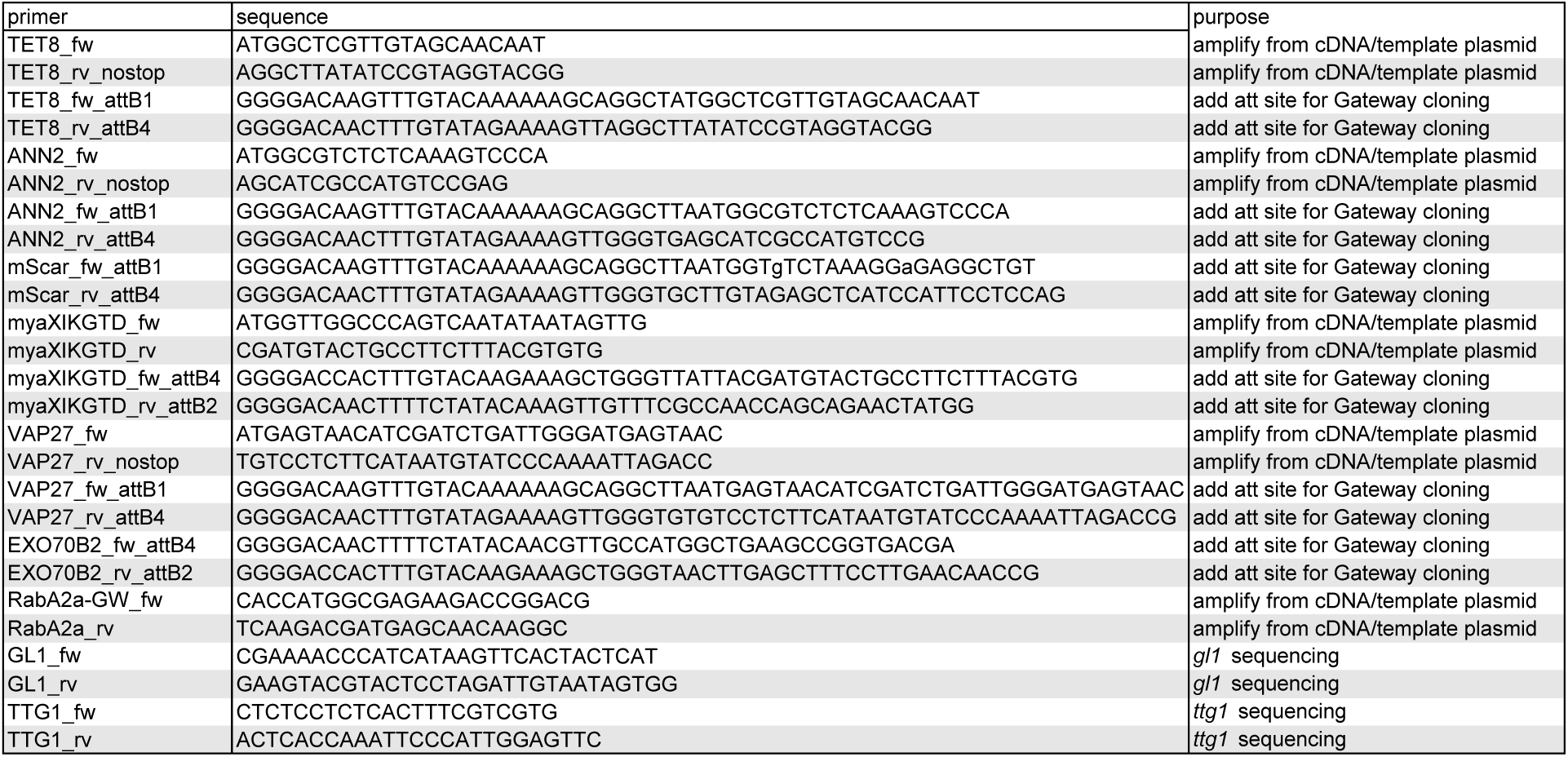
Primers used in this study.

### Transient expression

*Agrobacterium tumefaciens* strain GV3101 transformed with various expression plasmids was grown on solid media for 2 days prior to resuspension in 10 mM MgCl_2_ and 100 µM acetosyringone to a concentration of 0.2-0.6 OD depending on the level of protein expression of each plasmid. After incubation at room temperature for 2-3 h, bacteria were injected into leaves of 3-4 week old *N. benthamiana* plants grown in the same conditions as Arabidopsis but with 16-hour days. For dexamethasone (DEX) induction, a 50 µM solution of DEX was sprayed onto leaves 40 hours post injection. For co-immunoprecipitations, tissue was flash-frozen 8 h after DEX induction. For proximity labeling, biotin was injected into *N. benthamiana* leaves 4-6 hours post DEX, and tissue was flash frozen 3-8 h post biotin injection, depending on the levels of biotinylation observed over time for each construct. For confocal fluorescence microscopy, DEX was applied 48 h post injection and localization of fluorescence proteins was observed 16 h post DEX.

### Fungal infection

*Colletotrichum higginsianum* (*Ch*) isolate IMI349063A was grown from spore glycerol stock in 250 mL Erlenmeyer flasks capped with a foam plug on 100 mL of Mathur’s media (15 mM glucose, 0.15% w/v peptone, 0.05% w/v yeast extract, 5 mM magnesium sulfate heptahydrate, 20 mM potassium dihydrogen phosphate, pH 5.5 with KOH, 2% w/v agar) in the same room as Arabidopsis plants. After 10-14 days of growth, spores were harvested in distilled water by gently scrubbing the surface of culture vessels with a glass hook. Spores were washed by centrifuging at 500 × *g* for 5 min at 25°C twice. Spores were resuspended in 1 mL distilled water and concentration was determined using a hemocytometer.

For inoculation, 2 *µ*L of spore solution was placed onto each leaf of 1-week old Arabidopsis cotyledons germinated horizontally on ½ MS media, and spore solution was allowed to evaporate for 30 min in a sterile environment before placing in a humid chamber (100% relative humidity) in the plant growth conditions described above for 54 h until trypan blue staining.

For trypan blue staining of fungal structures, infected seedlings were placed in trypan blue solution (0.1% w/v trypan blue, 25% v/v lactic acid, 25% v/v phenol [TE buffer equilibrated pH 7.5], 25% v/v glycerol) in 2 mL microfuge tubes, caps were sealed, and tubes were incubated in boiling water for 1 min. Trypan blue solution was removed after overnight incubation, replaced with 95% ethanol, then replaced with 2.5 g/mL chloral hydrate solution for at least 16 h to destain. Cotyledons were mounted on slides in 50% v/v glycerol and observed using 5x, 20x, and 40x objectives (Invitrogen EVOS XL Core #AMEX1200). Images were captured at 40X from random locations on each cotyledon, and the infection stage of each appressoria was categorized in comparison to an infection index (Koch et al. 2025).

### Mass spectrometry (MS)

For protein digestion, samples containing pelleted density gradient fractions were denatured in 8 M urea in 100 mM ammonium bicarbonate. Samples were incubated for 45 min at 57°C with 10 mM Tris(2-carboxyethyl)phosphine hydrochloride to reduce cysteine residue side chains. These side chains were then alkylated with 20 mM iodoacetamide for 1 h in the dark at 21°C. The urea was diluted to 1 M urea using 100 mM ammonium bicarbonate. A total of 0.4 µg trypsin (Promega) was added, and the samples were digested for 14 h at 37°C. For MS, the resulting peptide solution was desalted using ZipTip pipette tips (EMD Millipore), dried down and resuspended in 0.1% formic acid. Peptides were analyzed by LC-MS on an Orbitrap Fusion Lumos equipped with an Easy NanoLC1200. Buffer A was 0.1% formic acid in water. Buffer B was 0.1% formic acid in 80% acetonitrile. Peptides were separated on a 90-min gradient from 0% B to 35% B. Peptides were fragmented by HCD at a relative collision energy of 32%. Precursor ions were measured in the Orbitrap with a resolution of 60,000. Fragment ions were measured in the Orbitrap with a resolution of 15,000. Data were analyzed using Proteome Discoverer (2.5) to interpret and quantify the relative amounts in a label free quantification manner. Data were searched against the Arabidopsis TAIR10 proteome downloaded on 2/12/2015. Trypsin was set as the protease with up to two missed cleavages allowed. Carbamidomethylation of cysteine residues was set as a fixed modification. Oxidation of methionine and protein N-terminal acetylation were set as variable modifications. A precursor mass tolerance of 10 ppm and a fragment ion quantification tolerance of 0.04 Da were used. Data were quantified using the Minora feature detector node within Proteome Discoverer. Proteins with either less than 2 peptides or less than 4% sequence coverage were eliminated from analysis. Protein intensity values were normalized by quantile normalization. Heatmaps for enrichment were produced in R using the ComplexHeatmap package, hierarchical clustering was done in R using Ward.D2 model, and figures were formatted in Adobe Illlustrator.

### Accession numbers

Genes and protein names used in this study are *Nicotiana benthamiana* PEN1 ortholog (Nbe.v1.1.chr15g41000), *Nicotiana benthamiana* RIN4 ortholog (Nbe.v1.1.chr06g07530), PEN1/SYP121 (AT3G11820), RIN4 (AT3G25070), TET8 (AT2G23810), ANN2 (AT5G65020), VAP27-1 or VAP27 (AT3G60600), VAP27-3 (At2G45140), RABA2a (At1g09630), SCD1 (AT1G49040), PATL1 (AT1G72150), RBCL/Rubisco Large Subunit (ATCG00490), γ2-COPI (AT4G34450), EXO70B2 (AT1G07000), AvrPphB (Q52430), Myosin XI-K (AT5g20490), ATG8a (At4g21980), ATG5 (AT5G17290), ATG2 (AT3G19190), ATG7 (At5g45900), SID2 (AT1G74710), ARA6 (AT1G24560), GL1 (AT3G27920), TTG1 (At5g24520), RPS2 (At4g26090), RPM1 (AT3G07040), PEN3 (AT1G59870), EXO70A1 (AT5G03540), EXO70B1 (AT5G58430), EXO70E1 (AT3G29400), EXO70E2 (AT5G61010), EXO70H1 (AT3G55150), CPC1 (AT2G46410), TRY82 (AT5G53200), TET9 (AT4G30430), CD63 (P08962), FREE1 (At1g20110), ALIX (AT1G15130). Additional accession numbers are included in Supplementary Datasets S1 and S2, which show mass spectrometry analysis of biotinylated proteins following EV protein proximity labeling in *N. benthamiana* (S1) and of P40 pellets isolated from autophagy mutants (S2).

## Acknowledgements

We are grateful for the assistance of Andras Kun and the IU Light Microscopy Imaging Center, Giovanni Gonzalez-Gutierrez and the IU Physical and Biochemical Instrumentation Facility, Barry Stein and the IU Electron Microscopy Center, and for equipment provided by the IU Biological Mass Spectrometry Center. We thank Jeff Dangl and Sebastian Y. Bednarek for providing seed of the *rin4rps2rpm1* and *scd1* Arabidopsis mutants and the Arabidopsis Biological Resource Center at The Ohio State University for providing all other Arabidopsis mutant lines used in this study. We are also indebted to Craig Pikaard for access to centrifuges. We also thank Chris Staiger for sharing cloning materials. We thank Melissa Blunck, Indiana University Undergraduate Research (IUUR), and the IUUR STEM Summer Research Program for supporting this research project and bringing us together.

## Author contributions

B.L.K. and R.W.I. designed the research; B.L.K., D.G., H.S., R.B., L.A, J.F., M.L.B., and B.R. performed the research; R.P. and J.T. performed mass spectrometry analyses, B.L.K. and R.W.I. analyzed the data; B.L.K. wrote the paper. R.W.I. edited the paper. All authors read and approved the manuscript before submission.

## Supplementary data

**Supplementary Dataset S1.** Mass spectrometry of biotinylated proteins following EV protein proximity labeling in *N. benthamiana*.

**Supplementary Dataset S2.** Mass spectrometry of proteins identified in EV pellet (P40) of *atg*- and *sid-*related Arabidopsis mutants.

## References

An Q, Ehlers K, Kogel KH, van Bel AJ, Huckelhoven R. Multivesicular compartments proliferate in susceptible and resistant *MLA12*-barley leaves in response to infection by the biotrophic powdery mildew fungus. New Phytol. 2006:172(3):563–576. 10.1111/j.1469-8137.2006.01844.x

An Q, Huckelhoven R, Kogel KH, van Bel AJ. Multivesicular bodies participate in a cell wall-associated defence response in barley leaves attacked by the pathogenic powdery mildew fungus. Cell Microbiol. 2006:8(6):1009–1019. 10.1111/j.1462-5822.2006.00683.x

Assaad FF, Qiu JL, Youngs H, Ehrhardt D, Zimmerli L, Kalde M, Wanner G, Peck SC, Edwards H, Ramonell K, Somerville CR, Thordal-Christensen H. The PEN1 syntaxin defines a novel cellular compartment upon fungal attack and is required for the timely assembly of papillae. Mol Biol Cell. 2004:15(11):5118–5129. 10.1091/mbc.e04-02-0140

Babst M. The Vps4p AAA ATPase regulates membrane association of a Vps protein complex required for normal endosome function. The EMBO Journal. 1998:17(11):2982–2993. 10.1093/emboj/17.11.2982

Babst M, Katzmann DJ, Snyder WB, Wendland B, Emr SD. Endosome-associated complex, ESCRT-II, recruits transport machinery for protein sorting at the multivesicular body. Developmental Cell. 2002:3(2):283–289. 10.1016/s1534-5807(02)00219-8

Bache KG, Brech A, Mehlum A, Stenmark H. Hrs regulates multivesicular body formation via ESCRT recruitment to endosomes. The Journal of Cell Biology. 2003:162(3):435–442. 10.1083/jcb.200302131

Baietti MF, Zhang Z, Mortier E, Melchior A, Degeest G, Geeraerts A, Ivarsson Y, Depoortere F, Coomans C, Vermeiren E, Zimmermann P, David G. Syndecan–syntenin–ALIX regulates the biogenesis of exosomes. Nature Cell Biology. 2012:14(7):677–685. 10.1038/ncb2502

Balkunde R, Deneer A, Bechtel H, Zhang B, Herberth S, Pesch M, Jaegle B, Fleck C, Hülskamp M. Identification of the trichome patterning core network using data from weak *ttg1* alleles to constrain the model space. Cell Reports. 2020:33(11):10.1016/j.celrep.2020.108497

Barman B, Ramirez M, Dawson TR, Liu Q, Weaver AM. Analysis of small EV proteomes reveals unique functional protein networks regulated by VAP-A. Proteomics. 2024:24(11):e2300099. 10.1002/pmic.202300099

Barman B, Sung BH, Krystofiak E, Ping J, Ramirez M, Millis B, Allen R, Prasad N, Chetyrkin S, Calcutt MW, Vickers K, Patton JG, Liu Q, Weaver AM. VAP-A and its binding partner CERT drive biogenesis of RNA-containing extracellular vesicles at ER membrane contact sites. Dev Cell. 2022:57(8):974–994 e978. 10.1016/j.devcel.2022.03.012

Beer KB, Rivas-Castillo J, Kuhn K, Fazeli G, Karmann B, Nance JF, Stigloher C, Wehman AM. Extracellular vesicle budding is inhibited by redundant regulators of TAT-5 flippase localization and phospholipid asymmetry. Proceedings of the National Academy of Sciences. 2018:115(6):10.1073/pnas.1714085115

Biller SJ, Schubotz F, Roggensack SE, Thompson AW, Summons RE, Chisholm SW. Bacterial vesicles in marine ecosystems. Science. 2014:343(6167):183–186. 10.1126/science.1243457

Boavida LC, Qin P, Broz M, Becker JD, McCormick S. Arabidopsis tetraspanins are confined to discrete expression domains and cell types in reproductive tissues and form homo- and heterodimers when expressed in yeast. Plant Physiol. 2013:163(2):696–712. 10.1104/pp.113.216598

Bobrie A, Colombo M, Krumeich S, Raposo G, Thery C. Diverse subpopulations of vesicles secreted by different intracellular mechanisms are present in exosome preparations obtained by differential ultracentrifugation. J Extracell Vesicles. 2012:1(10.3402/jev.v1i0.18397

Cai Q, Qiao L, Wang M, He B, Lin FM, Palmquist J, Huang SD, Jin H. Plants send small RNAs in extracellular vesicles to fungal pathogen to silence virulence genes. Science. 2018:360(6393):1126–1129. 10.1126/science.aar4142

Chaya T, Banerjee A, Rutter BD, Adekanye D, Ross J, Hu G, Innes RW, Caplan JL. The extracellular vesicle proteomes of *Sorghum bicolor* and *Arabidopsis thaliana* are partially conserved. Plant Physiol. 2024:194(3):1481–1497. 10.1093/plphys/kiad644

Chong YT, Gidda SK, Sanford C, Parkinson J, Mullen RT, Goring DR. Characterization of the *Arabidopsis thaliana* exocyst complex gene families by phylogenetic, expression profiling, and subcellular localization studies. New Phytol. 2010:185(2):401–419. 10.1111/j.1469-8137.2009.03070.x

Chow CM, Neto H, Foucart C, Moore I. Rab-A2 and Rab-A3 GTPases define a trans-golgi endosomal membrane domain in Arabidopsis that contributes substantially to the cell plate. Plant Cell. 2008:20(1):101–123. 10.1105/tpc.107.052001

Collins NC, Thordal-Christensen H, Lipka V, Bau S, Kombrink E, Qiu J-L, Hückelhoven R, Stein M, Freialdenhoven A, Somerville SC, Schulze-Lefert P. SNARE-protein-mediated disease resistance at the plant cell wall. Nature. 2003:425(6961):973–977. 10.1038/nature02076

Colombo M, Moita C, van Niel G, Kowal J, Vigneron J, Benaroch P, Manel N, Moita LF, Thery C, Raposo G. Analysis of ESCRT functions in exosome biogenesis, composition and secretion highlights the heterogeneity of extracellular vesicles. J Cell Sci. 2013:126(Pt 24):5553–5565. 10.1242/jcs.128868

Dagdas YF, Pandey P, Tumtas Y, Sanguankiattichai N, Belhaj K, Duggan C, Leary AY, Segretin ME, Contreras MP, Savage Z, Khandare VS, Kamoun S, Bozkurt TO. Host autophagy machinery is diverted to the pathogen interface to mediate focal defense responses against the Irish potato famine pathogen. Elife. 2018:7(10.7554/eLife.37476

Dinesh-Kumar SP, Nagalakshmi U, Wen Z, Yang X, Li Y, Zhang Y. TurboID-based proximity labeling for in planta identification of protein-protein interaction networks. Journal of Visualized Experiments. 2020:159):10.3791/60728-v

Ding Y, Wang J, Chun Lai JH, Ling Chan VH, Wang X, Cai Y, Tan X, Bao Y, Xia J, Robinson DG, Jiang L, Nakano A. Exo70E2 is essential for exocyst subunit recruitment and EXPO formation in both plants and animals. Molecular Biology of the Cell. 2014:25(3):412–426. 10.1091/mbc.e13-10-0586

Ebine K, Fujimoto M, Okatani Y, Nishiyama T, Goh T, Ito E, Dainobu T, Nishitani A, Uemura T, Sato MH, Thordal-Christensen H, Tsutsumi N, Nakano A, Ueda T. A membrane trafficking pathway regulated by the plant-specific RAB GTPase ARA6. Nat Cell Biol. 2011:13(7):853–859. 10.1038/ncb2270

Ehrlich MA, Schafer JF, Ehrlich HG. Lomasomes in wheat leaves infected by *Puccinia graminis* and *P. recondita*. Canadian Journal of Botany. 1968:46(1):17–20. 10.1139/b68-004

Erdmann S, Tschitschko B, Zhong L, Raftery MJ, Cavicchioli R. A plasmid from an Antarctic haloarchaeon uses specialized membrane vesicles to disseminate and infect plasmid-free cells. Nat Microbiol. 2017:2(10):1446–1455. 10.1038/s41564-017-0009-2

Falbel TG, Koch LM, Nadeau JA, Segui-Simarro JM, Sack FD, Bednarek SY. SCD1 is required for cytokinesis and polarized cell expansion in *Arabidopsis thaliana*. Development. 2003:130(17):4011–4024. 10.1242/dev.00619

Ghosh S, Innes RW. Expression of Arabidopsis extracellular vesicle protein markers in *Nicotiana benthamiana* reveals distinct vesicle subpopulations. bioRxiv. 2025:2025.2005.2006.652549. 10.1101/2025.05.06.652549

Gill S, Catchpole R, Forterre P. Extracellular membrane vesicles in the three domains of life and beyond. FEMS Microbiol Rev. 2019:43(3):273–303. 10.1093/femsre/fuy042

Gorlas A, Marguet E, Gill S, Geslin C, Guigner JM, Guyot F, Forterre P. Sulfur vesicles from Thermococcales: A possible role in sulfur detoxifying mechanisms. Biochimie. 2015:118(356–364. 10.1016/j.biochi.2015.07.026

Gu Y, Innes RW. The KEEP ON GOING Protein of Arabidopsis regulates intracellular protein trafficking and is degraded during fungal infection. The Plant Cell. 2012:24(11):4717–4730. 10.1105/tpc.112.105254

Guo H, Chitiprolu M, Roncevic L, Javalet C, Hemming FJ, Trung MT, Meng L, Latreille E, Tanese de Souza C, McCulloch D, Baldwin RM, Auer R, Côté J, Russell RC, Sadoul R, Gibbings D. Atg5 disassociates the V1V0-ATPase to promote exosome production and tumor metastasis independent of canonical macroautophagy. Developmental Cell. 2017:43(6):716–730.e717. 10.1016/j.devcel.2017.11.018

He B, Cai Q, Qiao L, Huang CY, Wang S, Miao W, Ha T, Wang Y, Jin H. RNA-binding proteins contribute to small RNA loading in plant extracellular vesicles. Nat Plants. 2021:7(3):342–352. 10.1038/s41477-021-00863-8

He Y, Xu J, Wang X, He X, Wang Y, Zhou J, Zhang S, Meng X. The Arabidopsis pleiotropic drug resistance transporters PEN3 and PDR12 mediate camalexin secretion for resistance to *Botrytis cinerea*. The Plant Cell. 2019:31(9):2206–2222. 10.1105/tpc.19.00239

Homma Y, Hiragi S, Fukuda M. Rab family of small GTPases: an updated view on their regulation and functions. The FEBS Journal. 2020:288(1):36–55. 10.1111/febs.15453

Huebbers JW, Büttgen K, Leissing F, Mantz M, Pauly M, Huesgen PF, Panstruga R. An advanced method for the release, enrichment and purification of high-quality *Arabidopsis thaliana* rosette leaf trichomes enables profound insights into the trichome proteome. Plant Methods. 2022:18(1):10.1186/s13007-021-00836-0

Huebbers JW, Caldarescu GA, Kubátová Z, Sabol P, Levecque SCJ, Kuhn H, Kulich I, Reinstädler A, Büttgen K, Manga-Robles A, Mélida H, Pauly M, Panstruga R, Žárský V. Interplay of EXO70 and MLO proteins modulates trichome cell wall composition and susceptibility to powdery mildew. The Plant Cell. 2024:36(4):1007–1035. 10.1093/plcell/koad319

Jeppesen DK, Fenix AM, Franklin JL, Higginbotham JN, Zhang Q, Zimmerman LJ, Liebler DC, Ping J, Liu Q, Evans R, Fissell WH, Patton JG, Rome LH, Burnette DT, Coffey RJ. Reassessment of exosome composition. Cell. 2019:177(2):428–445 e418. 10.1016/j.cell.2019.02.029

Jeppesen DK, Zhang Q, Franklin JL, Coffey RJ. Extracellular vesicles and nanoparticles: emerging complexities. Trends Cell Biol. 2023:33(8):667–681. 10.1016/j.tcb.2023.01.002

Jimenez-Jimenez S, Santana O, Lara-Rojas F, Arthikala MK, Armada E, Hashimoto K, Kuchitsu K, Salgado S, Aguirre J, Quinto C, Cardenas L. Differential tetraspanin genes expression and subcellular localization during mutualistic interactions in *Phaseolus vulgaris*. PLoS One. 2019:14(8):e0219765. 10.1371/journal.pone.0219765

Johnston EL, Zavan L, Bitto NJ, Petrovski S, Hill AF, Kaparakis-Liaskos M. Planktonic and biofilm-derived *Pseudomonas aeruginosa* outer membrane vesicles facilitate horizontal gene transfer of plasmid DNA. Microbiol Spectr. 2023:11(2):e0517922. 10.1128/spectrum.05179-22

Katzmann DJ, Babst M, Emr SD. Ubiquitin-dependent sorting into the multivesicular body pathway requires the function of a conserved endosomal protein sorting complex, ESCRT-I. Cell. 2001:106(2):145–155. 10.1016/s0092-8674(01)00434-2

Koch BL, Rutter BD, Borniego ML, Singla-Rastogi M, Gardner DM, Innes RW. Arabidopsis produces distinct subpopulations of extracellular vesicles that respond differentially to biotic stress, altering growth and infectivity of a fungal pathogen. J Extracell Vesicles. 2025:14(5):e70090. 10.1002/jev2.70090

Kurotani KI, Hirakawa H, Shirasawa K, Tagiri K, Mori M, Ramadan A, Ichihashi Y, Suzuki T, Tanizawa Y, An J, Winefield C, Waterhouse PM, Miura K, Nakamura Y, Isobe S, Notaguchi M. Establishing a comprehensive web-based analysis platform for *Nicotiana benthamiana* genome and transcriptome. Plant J. 2025:121(1):e17178. 10.1111/tpj.17178

Kurth EG, Peremyslov VV, Turner HL, Makarova KS, Iranzo J, Mekhedov SL, Koonin EV, Dolja VV. Myosin-driven transport network in plants. Proceedings of the National Academy of Sciences. 2017:114(8):10.1073/pnas.1620577114

Larios J, Mercier V, Roux A, Gruenberg J. ALIX- and ESCRT-III–dependent sorting of tetraspanins to exosomes. Journal of Cell Biology. 2020:219(3):10.1083/jcb.201904113

Lee Y, Rubio MC, Alassimone J, Geldner N. A mechanism for localized lignin deposition in the endodermis. Cell. 2013:153(2):402–412. 10.1016/j.cell.2013.02.045

Leidal AM, Huang HH, Marsh T, Solvik T, Zhang D, Ye J, Kai F, Goldsmith J, Liu JY, Huang YH, Monkkonen T, Vlahakis A, Huang EJ, Goodarzi H, Yu L, Wiita AP, Debnath J. The LC3-conjugation machinery specifies the loading of RNA-binding proteins into extracellular vesicles. Nat Cell Biol. 2020:22(2):187–199. 10.1038/s41556-019-0450-y

Li S, van Os GM, Ren S, Yu D, Ketelaar T, Emons AM, Liu CM. Expression and functional analyses of EXO70 genes in Arabidopsis implicate their roles in regulating cell type-specific exocytosis. Plant Physiol. 2010:154(4):1819–1830. 10.1104/pp.110.164178

Lin Y, Ding Y, Wang J, Kung C-H, Zhuang X, Yin Z, Xia Y, Robinson DG, Shen J, Lin H-X, Cui Y, Jiang L. EXPO and autophagosomes are distinct organelles in plants. Plant Physiology. 2015:10.1104/pp.15.00953

Liu D-A, Tao K, Wu B, Yu Z, Szczepaniak M, Rames M, Yang C, Svitkina T, Zhu Y, Xu F, Nan X, Guo W. A phosphoinositide switch mediates exocyst recruitment to multivesicular endosomes for exosome secretion. Nature Communications. 2023:14(1):10.1038/s41467-023-42661-0

Liu J, Cvirkaite-Krupovic V, Commere P-H, Yang Y, Zhou F, Forterre P, Shen Y, Krupovic M. Archaeal extracellular vesicles are produced in an ESCRT-dependent manner and promote gene transfer and nutrient cycling in extreme environments. The ISME Journal. 2021:15(10):2892–2905. 10.1038/s41396-021-00984-0

Liu N, Hou L, Chen X, Bao J, Chen F, Cai W, Zhu H, Wang L, Chen X. Arabidopsis TETRASPANIN8 mediates exosome secretion and glycosyl inositol phosphoceramide sorting and trafficking. Plant Cell. 2024:36(3):626–641. 10.1093/plcell/koad285

Liu N-J, Wang N, Bao J-J, Zhu H-X, Wang L-J, Chen X-Y. Lipidomic analysis reveals the importance of GIPCs in Arabidopsis leaf extracellular vesicles. Molecular Plant. 2020:13(10):1523–1532. 10.1016/j.molp.2020.07.016

Liu X-M, Ma L, Schekman R. Selective sorting of microRNAs into exosomes by phase-separated YBX1 condensates. eLife. 2021:10(10.7554/eLife.71982

Lu Q, Hope LW, Brasch M, Reinhard C, Cohen SN. TSG101 interaction with HRS mediates endosomal trafficking and receptor down-regulation. Proceedings of the National Academy of Sciences. 2003:100(13):7626–7631. 10.1073/pnas.0932599100

Mackey D, Belkhadir Y, Alonso JM, Ecker JR, Dangl JL. Arabidopsis RIN4 Is a target of the type III virulence effector AvrRpt2 and modulates RPS2-mediated resistance. Cell. 2003:112(3):379–389. 10.1016/s0092-8674(03)00040-0

Mackey D, Holt BF, 3rd, Wiig A, Dangl JL. RIN4 interacts with *Pseudomonas syringae* type III effector molecules and is required for RPM1-mediated resistance in Arabidopsis. Cell. 2002:108(6):743–754. 10.1016/s0092-8674(02)00661-x

Margets A, Foster J, Kumar A, Maier TR, Masonbrink R, Mejias J, Baum TJ, Innes RW. The soybean cyst nematode effector Cysteine Protease 1 (CPR1) targets a mitochondrial soybean branched-chain amino acid aminotransferase (GmBCAT1). Molecular Plant-Microbe Interactions®. 2024:37(11):751–764. 10.1094/mpmi-06-24-0068-r

Markovic V, Kulich I, Zarsky V. Functional specialization within the EXO70 gene family in Arabidopsis. Int J Mol Sci. 2021:22(14):10.3390/ijms22147595

Matern A, Böttcher C, Eschen-Lippold L, Westermann B, Smolka U, Döll S, Trempel F, Aryal B, Scheel D, Geisler M, Rosahl S. A substrate of the ABC transporter PEN3 stimulates bacterial flagellin (flg22)-induced callose deposition in *Arabidopsis thaliana*. Journal of Biological Chemistry. 2019:294(17):6857–6870. 10.1074/jbc.RA119.007676

Matsumura M, Nomoto M, Itaya T, Aratani Y, Iwamoto M, Matsuura T, Hayashi Y, Mori T, Skelly MJ, Yamamoto YY, Kinoshita T, Mori IC, Suzuki T, Betsuyaku S, Spoel SH, Toyota M, Tada Y. Mechanosensory trichome cells evoke a mechanical stimuli–induced immune response in *Arabidopsis thaliana*. Nature Communications. 2022:13(1):10.1038/s41467-022-28813-8

Mayers JR, Hu T, Wang C, Cardenas JJ, Tan Y, Pan J, Bednarek SY. SCD1 and SCD2 form a complex that functions with the exocyst and RabE1 in exocytosis and cytokinesis. Plant Cell. 2017:29(10):2610–2625. 10.1105/tpc.17.00409

Mei K, Li Y, Wang S, Shao G, Wang J, Ding Y, Luo G, Yue P, Liu J-J, Wang X, Dong M-Q, Wang H-W, Guo W. Cryo-EM structure of the exocyst complex. Nature Structural & Molecular Biology. 2018:25(2):139–146. 10.1038/s41594-017-0016-2

Melotto M, Underwood W, Koczan J, Nomura K, He SY. Plant stomata function in innate immunity against bacterial invasion. Cell. 2006:126(5):969–980. 10.1016/j.cell.2006.06.054

Meyer D, Pajonk S, Micali C, O’Connell R, Schulze-Lefert P. Extracellular transport and integration of plant secretory proteins into pathogen-induced cell wall compartments. Plant Journal. 2009:57(6):986–999. 10.1111/j.1365-313X.2008.03743.x

Micali CO, Neumann U, Grunewald D, Panstruga R, O’Connell R. Biogenesis of a specialized plant-fungal interface during host cell internalization of *Golovinomyces orontii* haustoria. Cell Microbiol. 2011:13(2):210–226. 10.1111/j.1462-5822.2010.01530.x

Mills J, Gebhard LJ, Schubotz F, Shevchenko A, Speth DR, Liao Y, Duggin IG, Marchfelder A, Erdmann S. Extracellular vesicle formation in Euryarchaeota is driven by a small GTPase. Proc Natl Acad Sci U S A. 2024:121(10):e2311321121. 10.1073/pnas.2311321121

Mosesso N, Lerner NS, Bläske T, Groh F, Maguire S, Niedermeier ML, Landwehr E, Vogel K, Meergans K, Nagel M-K, Drescher M, Stengel F, Hauser K, Isono E. Arabidopsis CaLB1 undergoes phase separation with the ESCRT protein ALIX and modulates autophagosome maturation. Nature Communications. 2024:15(1):10.1038/s41467-024-49485-6

Muzioł T, Pineda-Molina E, Ravelli RB, Zamborlini A, Usami Y, Göttlinger H, Weissenhorn W. Structural basis for budding by the ESCRT-III factor CHMP3. Developmental Cell. 2006:10(6):821–830. 10.1016/j.devcel.2006.03.013

Nabhan JF, Hu R, Oh RS, Cohen SN, Lu Q. Formation and release of arrestin domain-containing protein 1-mediated microvesicles (ARMMs) at plasma membrane by recruitment of TSG101 protein. Proceedings of the National Academy of Sciences. 2012:109(11):4146–4151. 10.1073/pnas.1200448109

Nager AR, Goldstein JS, Herranz-Pérez V, Portran D, Ye F, Garcia-Verdugo JM, Nachury MV. An actin network dispatches ciliary GPCRs into extracellular vesicles to modulate sgnaling. Cell. 2017:168(1-2):252–263.e214. 10.1016/j.cell.2016.11.036

Nawrath C, Metraux JP. Salicylic acid induction-deficient mutants of Arabidopsis express *PR-2* and *PR-5* and accumulate high levels of camalexin after pathogen inoculation. Plant Cell. 1999:11(8):1393–1404. 10.1105/tpc.11.8.1393

Nielsen ME, Feechan A, Bohlenius H, Ueda T, Thordal-Christensen H. Arabidopsis ARF-GTP exchange factor, GNOM, mediates transport required for innate immunity and focal accumulation of syntaxin PEN1. Proc Natl Acad Sci U S A. 2012:109(28):11443–11448. 10.1073/pnas.1117596109

Oliveira DL, Nakayasu ES, Joffe LS, Guimaraes AJ, Sobreira TJ, Nosanchuk JD, Cordero RJ, Frases S, Casadevall A, Almeida IC, Nimrichter L, Rodrigues ML. Characterization of yeast extracellular vesicles: evidence for the participation of different pathways of cellular traffic in vesicle biogenesis. PLoS One. 2010:5(6):e11113. 10.1371/journal.pone.0011113

Ortmannová J, Sekereš J, Kulich I, Šantrůček J, Dobrev P, Žárský V, Pečenková T, Monaghan J. Arabidopsis EXO70B2 exocyst subunit contributes to papillae and encasement formation in antifungal defence. Journal of Experimental Botany. 2022:73(3):742–755. 10.1093/jxb/erab457

Pandey P, Leary AY, Tumtas Y, Savage Z, Dagvadorj B, Duggan C, Yuen ELH, Sanguankiattichai N, Tan E, Khandare V, Connerton AJ, Yunusov T, Madalinski M, Mirkin FG, Schornack S, Dagdas Y, Kamoun S, Bozkurt TO. An oomycete effector subverts host vesicle trafficking to channel starvation-induced autophagy to the pathogen interface. eLife. 2021:10(10.7554/eLife.65285

Pang L, Ma Z, Zhang X, Huang Y, Li R, Miao Y, Li R. The small GTPase RABA2a recruits SNARE proteins to regulate the secretory pathway in parallel with the exocyst complex in Arabidopsis. Mol Plant. 2022:15(3):398–418. 10.1016/j.molp.2021.11.008

Pečenková T, Hála M, Kulich I, Kocourková D, Drdová E, Fendrych M, Toupalová H, Žárský V. The role for the exocyst complex subunits Exo70B2 and Exo70H1 in the plant–pathogen interaction. Journal of Experimental Botany. 2011:62(6):2107–2116. 10.1093/jxb/erq402

Peng X, Yang L, Ma Y, Li X, Yang S, Li Y, Wu B, Tang S, Zhang F, Zhang B, Liu J, Li H. IKKβ activation promotes amphisome formation and extracellular vesicle secretion in tumor cells. Biochimica et Biophysica Acta (BBA) - Molecular Cell Research. 2021:1868(1):10.1016/j.bbamcr.2020.118857

Pereira C, Stalder D, Anderson GSF, Shun-Shion AS, Houghton J, Antrobus R, Chapman MA, Fazakerley DJ, Gershlick DC. The exocyst complex is an essential component of the mammalian constitutive secretory pathway. Journal of Cell Biology. 2023:222(5):10.1083/jcb.202205137

Qi D, DeYoung BJ, Innes RW. Structure-function analysis of the coiled-coil and leucine-rich repeat domains of the RPS5 disease resistance protein. Plant Physiol. 2012:158(4):1819–1832. 10.1104/pp.112.194035

Qin L, Liu L, Tu J, Yang G, Wang S, Quilichini TD, Gao P, Wang H, Peng G, Blancaflor EB, Datla R, Xiang D, Wilson KE, Wei Y. The ARP2/3 complex, acting cooperatively with Class I formins, modulates penetration resistance in Arabidopsis against powdery mildew invasion. Plant Cell. 2021:33(9):3151–3175. 10.1093/plcell/koab170

Raiborg C, Bache KG, Gillooly DJ, Madshus IH, Stang E, Stenmark H. Hrs sorts ubiquitinated proteins into clathrin-coated microdomains of early endosomes. Nature Cell Biology. 2002:4(5):394–398. 10.1038/ncb791

Raiborg C, Stenmark H. The ESCRT machinery in endosomal sorting of ubiquitylated membrane proteins. Nature. 2009:458(7237):445–452. 10.1038/nature07961

Raposo G, Nijman HW, Stoorvogel W, Liejendekker R, Harding CV, Melief CJ, Geuze HJ. B lymphocytes secrete antigen-presenting vesicles. J Exp Med. 1996:183(3):1161–1172. 10.1084/jem.183.3.1161

Redditt TJ, Chung EH, Karimi HZ, Rodibaugh N, Zhang Y, Trinidad JC, Kim JH, Zhou Q, Shen M, Dangl JL, Mackey D, Innes RW. AvrRpm1 functions as an ADP-ribosyl transferase to modify NOI Domain-containing proteins, including Arabidopsis and soybean RPM1-Interacting Protein4. Plant Cell. 2019:31(11):2664–2681. 10.1105/tpc.19.00020R2

Regente M, Pinedo M, San Clemente H, Balliau T, Jamet E, de la Canal L. Plant extracellular vesicles are incorporated by a fungal pathogen and inhibit its growth. J Exp Bot. 2017:68(20):5485–5495. 10.1093/jxb/erx355

Regmi KC, Ghosh S, Koch B, Neumann U, Stein B, O’Connell RJ, Innes RW. Three-dimensional ultrastructure of Arabidopsis cotyledons infected with *Colletotrichum higginsianum*. Mol Plant Microbe Interact. 2024:37(4):396–406. 10.1094/MPMI-05-23-0068-R

Rubiato HM, Liu M, O’Connell RJ, Nielsen ME. Plant SYP12 syntaxins mediate an evolutionarily conserved general immunity to filamentous pathogens. Elife. 2022:11(10.7554/eLife.73487

Russell MRG, Shideler T, Nickerson DP, West M, Odorizzi G. Class E compartments form in response to ESCRT dysfunction in yeast due to hyperactivity of the Vps21 Rab GTPase. Journal of Cell Science. 2012:10.1242/jcs.111310

Rutter BD, Innes RW. Extracellular vesicles isolated from the leaf apoplast carry stress-response proteins. Plant Physiol. 2017:173(1):728–741. 10.1104/pp.16.01253

Schuurink R, Tissier A. Glandular trichomes: micro-organs with model status? New Phytologist. 2019:225(6):2251–2266. 10.1111/nph.16283

Speth EB, Imboden L, Hauck P, He SY. Subcellular localization and functional analysis of the Arabidopsis GTPase RabE Plant Physiology. 2009:149(4):1824–1837. 10.1104/pp.108.132092

Stefano G, Renna L, Wormsbaecher C, Gamble J, Zienkiewicz K, Brandizzi F. Plant endocytosis requires the ER membrane-anchored proteins VAP27-1 and VAP27-3. Cell Reports. 2018:23(8):2299–2307. 10.1016/j.celrep.2018.04.091

Stein M, Dittgen J, Sanchez-Rodriguez C, Hou BH, Molina A, Schulze-Lefert P, Lipka V, Somerville S. Arabidopsis PEN3/PDR8, an ATP binding cassette transporter, contributes to nonhost resistance to inappropriate pathogens that enter by direct penetration. Plant Cell. 2006:18(3):731–746. 10.1105/tpc.105.038372

Stuffers S, Sem Wegner C, Stenmark H, Brech A. Multivesicular endosome biogenesis in the absence of ESCRTs. Traffic. 2009:10(7):925–937. 10.1111/j.1600-0854.2009.00920.x

Tapken W, Murphy AS. Membrane nanodomains in plants: capturing form, function, and movement. Journal of Experimental Botany. 2015:66(6):1573–1586. 10.1093/jxb/erv054

Temoche-Diaz MM, Shurtleff MJ, Nottingham RM, Yao J, Fadadu RP, Lambowitz AM, Schekman R. Distinct mechanisms of microRNA sorting into cancer cell-derived extracellular vesicle subtypes. Elife. 2019:8(10.7554/eLife.47544

Tkach M, Kowal J, Zucchetti AE, Enserink L, Jouve M, Lankar D, Saitakis M, Martin-Jaular L, Thery C. Qualitative differences in T-cell activation by dendritic cell-derived extracellular vesicle subtypes. EMBO J. 2017:36(20):3012–3028. 10.15252/embj.201696003

Trajkovic K, Hsu C, Chiantia S, Rajendran L, Wenzel D, Wieland F, Schwille P, Brugger B, Simons M. Ceramide triggers budding of exosome vesicles into multivesicular endosomes. Science. 2008:319(5867):1244-1247. 10.1126/science.1153124

Tripathy MK, Deswal R, Sopory SK. Plant RABs: role in development and in abiotic and biotic stress responses. Current Genomics. 2021:22(1):26–40. 10.2174/1389202922666210114102743

van der Hoorn RAL, Kamoun S. From guard to decoy: a new model for perception of plant pathogen effectors. The Plant Cell. 2008:20(8):2009–2017. 10.1105/tpc.108.060194

Verweij FJ, Revenu C, Arras G, Dingli F, Loew D, Pegtel DM, Follain G, Allio G, Goetz JG, Zimmermann P, Herbomel P, Del Bene F, Raposo G, van Niel G. Live tracking of inter-organ communication by endogenous exosomes in vivo. Dev Cell. 2019:48(4):573–589 e574. 10.1016/j.devcel.2019.01.004

Vinatzer BA, Teitzel GM, Lee MW, Jelenska J, Hotton S, Fairfax K, Jenrette J, Greenberg JT. The type III effector repertoire of *Pseudomonas syringae* pv. *syringae* B728a and its role in survival and disease on host and non-host plants. Mol Microbiol. 2006:62(1):26–44. 10.1111/j.1365-2958.2006.05350.x

Wang J, Ding Y, Wang J, Hillmer S, Miao Y, Lo SW, Wang X, Robinson DG, Jiang L. EXPO, an exocyst-positive organelle distinct from multivesicular endosomes and autophagosomes, mediates cytosol to cell wall exocytosis in Arabidopsis and tobacco cells The Plant Cell. 2010:22(12):4009–4030. 10.1105/tpc.110.080697

Wang P, Richardson C, Hawkins TJ, Sparkes I, Hawes C, Hussey PJ. Plant VAP27 proteins: domain characterization, intracellular localization and role in plant development. New Phytol. 2016:210(4):1311–1326. 10.1111/nph.13857

Wang S, He B, Wu H, Cai Q, Ramirez-Sanchez O, Abreu-Goodger C, Birch PRJ, Jin H. Plant mRNAs move into a fungal pathogen via extracellular vesicles to reduce infection. Cell Host Microbe. 2024:32(1):93–105 e106. 10.1016/j.chom.2023.11.020

Wang Y, Li S, Mokbel M, May AI, Liang Z, Zeng Y, Wang W, Zhang H, Yu F, Sporbeck K, Jiang L, Aland S, Agudo-Canalejo J, Knorr RL, Fang X. Biomolecular condensates mediate bending and scission of endosome membranes. Nature. 2024:634(8036):1204–1210. 10.1038/s41586-024-07990-0

Williams JK, Ngo JM, Lehman IM, Schekman R. Annexin A6 mediates calcium-dependent exosome secretion during plasma membrane repair. Elife. 2023:12(10.7554/eLife.86556

Willms E, Johansson HJ, Mager I, Lee Y, Blomberg KE, Sadik M, Alaarg A, Smith CI, Lehtio J, El Andaloussi S, Wood MJ, Vader P. Cells release subpopulations of exosomes with distinct molecular and biological properties. Sci Rep. 2016:6(22519. 10.1038/srep22519

Wu J, Liu Y. Stomata-pathogen interactions: over a century of research. Trends Plant Sci. 2022:27(10):964–967. 10.1016/j.tplants.2022.07.004

Yang L, Qin L, Liu G, Peremyslov VV, Dolja VV, Wei Y. Myosins XI modulate host cellular responses and penetration resistance to fungal pathogens. Proc Natl Acad Sci U S A. 2014:111(38):13996–14001. 10.1073/pnas.1405292111

Yoshimoto K, Jikumaru Y, Kamiya Y, Kusano M, Consonni C, Panstruga R, Ohsumi Y, Shirasu K. Autophagy negatively regulates cell death by controlling NPR1-dependent salicylic acid signaling during senescence and the innate immune response in Arabidopsis. Plant Cell. 2009:21(9):2914–2927. 10.1105/tpc.109.068635

Zarnowski R, Sanchez H, Covelli AS, Dominguez E, Jaromin A, Bernhardt J, Mitchell KF, Heiss C, Azadi P, Mitchell A, Andes DR. *Candida albicans* biofilm-induced vesicles confer drug resistance through matrix biogenesis. PLoS Biol. 2018:16(10):e2006872. 10.1371/journal.pbio.2006872

Zarsky V, Cvrckova F, Potocky M, Hala M. Exocytosis and cell polarity in plants - exocyst and recycling domains. New Phytol. 2009:183(2):255–272. 10.1111/j.1469-8137.2009.02880.x

Zeyen RJ, Bushnel WR. Papilla response of barley epidermal cells caused by *Erysiphe graminis*: rate and method of deposition determined by microcinematography and transmission electron microscopy. Canadian Journal of Botany. 1979:57(8):898–913. 10.1139/b79-111

Zhang J, Qiu Y, Xu K. Characterization of GFP-AtPEN1 as a marker protein for extracellular vesicles isolated from *Nicotiana benthamiana* leaves. Plant Signal Behav. 2020:15(9):1791519. 10.1080/15592324.2020.1791519

Zhang W, Huang L, Zhang C, Staiger CJ. Arabidopsis myosin XIK interacts with the exocyst complex to facilitate vesicle tethering during exocytosis. The Plant Cell. 2021:33(7):2454–2478. 10.1093/plcell/koab116

Zhao J, Bui MT, Ma J, Kunzl F, Picchianti L, De La Concepcion JC, Chen Y, Petsangouraki S, Mohseni A, Garcia-Leon M, Gomez MS, Giannini C, Gwennogan D, Kobylinska R, Clavel M, Schellmann S, Jaillais Y, Friml J, Kang BH, Dagdas Y. Plant autophagosomes mature into amphisomes prior to their delivery to the central vacuole. J Cell Biol. 2022:221(12):10.1083/jcb.202203139

Zhao K, Bleackley M, Chisanga D, Gangoda L, Fonseka P, Liem M, Kalra H, Al Saffar H, Keerthikumar S, Ang C-S, Adda CG, Jiang L, Yap K, Poon IK, Lock P, Bulone V, Anderson M, Mathivanan S. Extracellular vesicles secreted by *Saccharomyces cerevisiae* are involved in cell wall remodelling. Communications Biology. 2019:2(1):10.1038/s42003-019-0538-8

Zhu X, Wang W, Sun S, Chng CP, Xie Y, Zhu K, He D, Liang Q, Ma Z, Wu X, Zheng X, Gao W, Miserez A, Gao C, Yu J, Huang C, Groves JT, Miao Y. Bacterial XopR subverts RIN4 complex-mediated plant immunity via plasma membrane-associated percolation. Dev Cell. 2025:10.1016/j.devcel.2025.03.002

